# The *PIF1-MIR408-Plantacyanin* Repression Cascade Regulates Light Dependent Seed Germination

**DOI:** 10.1101/2020.07.20.212340

**Authors:** Anlong Jiang, Zhonglong Guo, Jiawei Pan, Yan Zhuang, Daqing Zuo, Chen Hao, Zhaoxu Gao, Peiyong Xin, Jinfang Chu, Shangwei Zhong, Lei Li

## Abstract

Light-sensing seed germination is a vital process for the seed plants. A decisive event in light-induced germination is degradation of the central repressor PHYTOCHROME INTERACTING FACTOR1 (PIF1). It is also known that the balance between gibberellic acid (GA) and abscisic acid (ABA) critically controls germination. But the cellular mechanisms linking PIF1 turnover to hormonal rebalancing remain elusive. Here, employing far-red light-induced *Arabidopsis* seed germination as the experimental system, we identified Plantacyanin (PLC) as an inhibitor of germination, which is a storage vacuole-associated blue copper protein highly expressed in mature seed and rapidly silenced during germination. Molecular analyses showed that PIF1 directly binds to the *MIR408* promoter and represses miR408 accumulation, which in turn post-transcriptionally modulates *PLC* abundance, thus forming the *PIF1-MIR408-PLC* repression cascade for translating PIF1 turnover to PLC turnover during early germination. Genetic analysis, RNA-sequencing, and hormone quantification revealed that *PLC* is necessary and sufficient to maintain the *PIF1*-mediated seed transcriptome and the low-GA-high-ABA state. Furthermore, we found that PLC domain organization and regulation by miR408 are conserved features in seed plants. These results unraveled a cellular mechanism whereby PIF1-relayed external light signals are converted through PLC-based copper mobilization into internal hormonal profiles for controlling seed germination.

## Introduction

The seed is an embryonic plant enclosed in a protective capsule. After reaching full size, the embryo undergoes elaborate dehydration to establish a dormant state that helps the embryo withstand extreme environments and survive for long periods (Bewley 1997; Finch-Savage and Leubner-Metzger, 2006; Angelovici et al., 2010). Given the right environmental conditions, the desiccated seed germinates by taking up water and resuming embryo development, utilizing energy and nutrient stored in the seed (Bewley 1997; Finch-Savage and Leubner-Metzger, 2006; Finkelstein et al., 2008; Née et al., 2017). GA and ABA are the main plant hormones that control seed dormancy and germination. As the embryo matures, ABA is synthesized and signals the embryo to initiate the buildup of storage compounds and undergo desiccation (Nambara and Marion-Poll, 2005; Finkelstein et al., 2008; Angelovici et al., 2010; Shu et al., 2016). ABA is also important for maintaining seed dormancy and preventing precocious germination (Bewley 1997; Finch-Savage and Leubner-Metzger, 2006; Finkelstein et al., 2008). Conversely, GA is a crucial hormone to break down dormancy and promote germination (Kallioo and Piiroinen 1959; Bewley 1997; Finch-Savage and Leubner-Metzger, 2006; Yamaguchi, 2008). It has been well-established that the GA/ABA balance critically determines the germination capacity (Nambara and Marion-Poll, 2005; Yamaguchi, 2008; Shu et al., 2016; Née et al., 2017).

Seed monitors a wide range of environmental factors, including ambient light, for germination decision-making (Oh et al., 2004; Finch-Savage and Leubner-Metzger, 2006; Seo et al., 2009; Jiang et al., 2016). Molecular mechanisms of light perception and signaling during germination are well understood in the model plant *Arabidopsis*. The basic helix-loop-helix type transcription factor PIF1 is an essential negative regulator of light-dependent germination (Oh et al., 2004; Leivar and Quail, 2010; Shi et al., 2015). PIF1 is stabilized by DE-ETIOLATED 1 and other signaling molecules and thus highly accumulates in the seed kept in darkness (Oh et al., 2004; 2006; Shi et al., 2015). Under light irradiation, phytochromes are activated and enter the nucleus to interact with PIF1, thereby reducing its activity and promoting its degradation via the 26S proteasome (Oh et al., 2006; Castillon et al., 2007; Shen et al., 2008; Leivar and Quail, 2010). Rapid removal of PIF1 is critical for maintaining the light-regulated transcriptome in imbibed seed (Oh et al., 2009; Shi et al., 2013; Pfeiffer et al., 2014) and ultimately establishing the high-GA-low-ABA state (Oh et al., 2006; 2007). However, extensive search has not revealed a direct link between PIF1 and genes involved in GA and ABA metabolism (Oh et al., 2007; 2009; Cho et al., 2012). Consequently, the cellular events ensued by PIF1 turnover that lead to hormonal rebalancing have not been elucidated and it is not known whether these events are conserved in seed plants.

A fundamental cellular process during seed germination is the mobilization of mineral nutrients sequestered in the storage vacuoles to sustain embryo growth before efficient uptake systems are established in the root (Lanquar et al., 2005; Kim et al., 2006; Roschzttardtz et al., 2009; Née et al., 2017; Paszkiewicz et al., 2017). Studies in *Arabidopsis* and other plants have shown that transition metals are released from vacuolar stores via tonoplast-localized transporters and then transported to various cellular destinations (Lanquar et al., 2005; Kim et al., 2006; Eroglu et al., 2017). While the physiological consequences of insufficient or mis-regulated mineral mobilization have been abundantly documented (Lanquar et al., 2005; Kim et al., 2006), whether metal mobilization contributes to hormonal profile establishment in light dependent germination is not well characterized.

Copper is an essential transition metal by serving as the cofactor for a number of cuproproteins with vital functions (Burkhead et al., 2009; Peñarrubia et al., 2015). Because free cellular copper is highly reactive and produces detrimental hydroxyl radicals, elaborate homeostasis and transport systems are present for the precise control of copper delivery to specific targets (Burkhead et al., 2009). The *Arabidopsis* genome encodes approximately 260 copper dependent proteins (Schulten et al., 2019). Among them, small blue copper proteins, containing a characteristic mononuclear type-I copper binding site, play important roles in redox reactions and copper homeostasis (Rydén and Hunt 1993; Guss et al., 1998; De Rienzo et al. 2000; Giri et al. 2004). Plastocyanin is the most abundant small blue copper protein in plants and an indispensable electron carrier in the Z-scheme of photosynthesis (Molina-Heredia et al., 2003; Weigel et al., 2003). Plants also have a specific family of blue copper proteins called phytocyanins that are divided into four subfamilies, including PLCs, uclacyanins, stellacyanins, and early nodulin-like proteins, based on differences in the copper binding site and domain organization (Guss et al., 1998; Nersissian et al., 1998; Sun et al., 2019). Phytocyanins have been widely implicated in plant development processes such as pollen tube chemotropism and nodule development (Kim et al. 2003; Dong et al. 2005; Sun et al., 2019). They have also been implicated in stress responses such as pathogen resistance and drought and salinity tolerance (Jung and Hwang 2000; Ruan et al. 2011; Feng et al., 2013). However, their involvement in seed germination has not been investigated.

In this study, we focused on *PLC* that is highly expressed in the seed with contrasting expression patterns during seed development and germination. Through comprehensive molecular and genetic analyses, we delineated the *PIF1*-*MIR408*-*PLC* repression cascade for regulating PLC turnover during far-red light induced germination. We showed that PLC locates to the storage vacuole and is necessary and sufficient to maintain *PIF1*-mediated seed transcriptome and the low-GA-high-ABA state. These results unraveled PLC-based copper mobilization as a potentially conserved cellular mechanism for converting PIF1-relayed light signals into hormonal profiles that control seed germination.

## RESULTS

### *PLC* Exhibits Distinctive Expression Pattern in Seed Development and Germination

To investigate whether the phytocyanin encoding genes are involved in seed germination, we examined their expression pattern using a gene expression atlas in *Arabidopsis* (eFP Browser; Winter et al., 2007). We found that 31 of the 37 phytocyanin genes were expressed during seed formation and germination (Figure 1A). Among these, the single gene encoding PLC (At2g02850) exhibited the highest expression level in mature seed (Figure 1A). *PLC* transcript level was low in early seed development, but was drastically induced at seed stage 7, progressively increased thereafter, and peaked at seed maturation (Figure 1B). Interestingly, *PLC* was drastically silenced upon vernalization-induced germination (Figure 1B). To validate PLC expression pattern in the germination phase at the protein level, we generated the *pPLC:PLC-GFP* transgenic *Arabidopsis* plants in which the *PLC* coding sequence was fused to the green fluorescence protein (GFP) and driven by the native *PLC* promoter. Immunoblotting with an antibody against GFP revealed that the level of the PLC-GFP fusion protein was drastically decreased 24 h after vernalization (Figure 1C). These results indicate that *PLC* is highly expressed in mature seed and silenced during germination.

**Figure 1.**
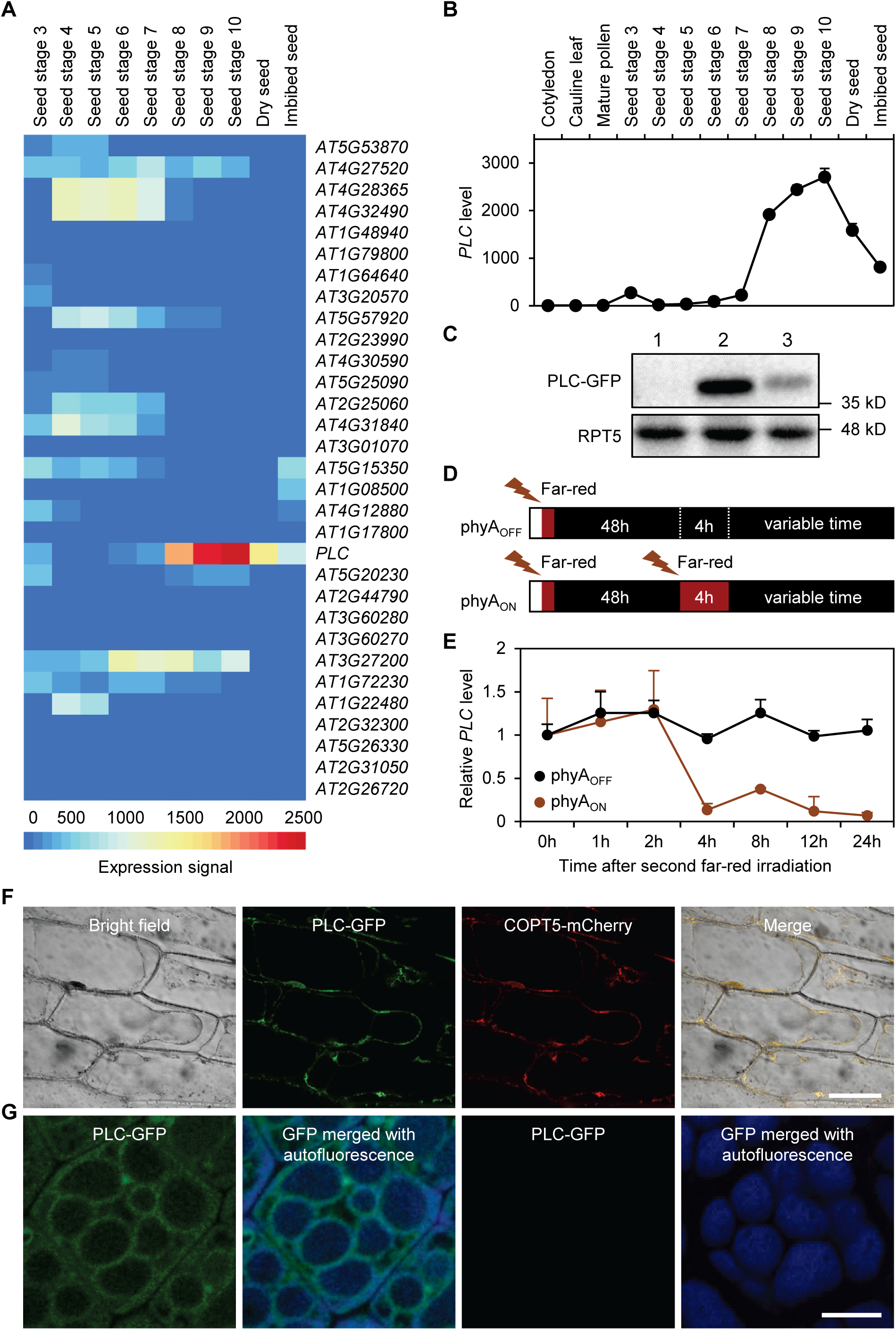
*PLC* Is Induced during Seed Development and Rapidly Silenced during Germination. (**A**) Global expression profile of 31 phytocyanin genes in the seed based on data in the *Arabidopsis* eFP Browser. (**B**) Comparison of *PLC* expression pattern in the seed and other organs, using data from the eFP Browser. (**C**) Detection of PLC-GFP in *pPLC:PLC-GFP* plants using an anti-GFP antibody. 1, seedling; 2, dry seed; 3, vernalized seed. Size markers are indicated on the right. RPT5 was used as the loading control. (**D**) Diagram illustrating the phyA_OFF_ and phyA_ON_ regimes in which imbibed seeds were sequentially treated with the indicated light conditions. (**E**) Relative *PLC* transcript level during the time course of phyA_OFF_ and phyA_ON_ determined by RT-qPCR analysis. Data are mean ± SD (n = 3). (**F**) Subcellular localization of PLC. PLC-GFP and COPT5-mCherry were transiently expressed in the same onion epidermal cells and examined by confocal fluorescence microscopy. (**G**) Co-localization of GFP fluorescence with vacuole autofluorescence in cotyledon cells of imbibed *pPLC:PLC-GFP* seed in phyA_OFF_ (left two panels) and phyA_ON_ (right two panels). Bars, 10 μm.

To elucidate the dynamics of *PLC* repression during light-induced germination, we employed a previously described far-red light initiated, phytochrome A (phyA) dependent germination assay (Oh et al., 2006; 2009; Cho et al., 2012). In the so-called phyA_OFF_ regime of this assay, imbibed seeds are exposed to brief far-red light to inactivate phyB, then kept in darkness to allow inactive phyA to accumulate that does not break dormancy. In the phyA_ON_ regime, a second far-red irradiation with longer duration is used to activate phyA that induces germination (Figure 1D). Using reverse transcription coupled quantitative PCR (RT-qPCR), we observed that *PLC* transcript level remained steady during the time course of phyA_OFF_ (Figure 1E). In contrast, after the second far-red irradiation in phyA_ON_, *PLC* level was maintained only until the 2 h time point, but drastically reduced by 4 h, and remained low thereafter (Figure 1E).

To reveal subcellular localization of PLC, we transiently expressed the PLC-GFP reporter in onion epidermal cells and found that the GFP signals predominantly aligned the periphery of the central vacuole (Figure 1F). COPT5 is a member of the CTR-like high-affinity Cu transporters residing on the tonoplast (Klaumann et al., 2011). We found that PLC-GFP colocalized with mCherry-tagged COPT5 co-expressed in the same onion epidermal cell (Figure 1F), indicating that transiently expressed PLC is associated with the vacuole. To examine PLC localization in the seed, we utilized the *pPLC:PLC-GFP* transgenic line. In imbibed seed kept in darkness, GFP fluorescence was observed surrounding autofluorescence of storage vacuoles (Figure 1G) (Paszkiewicz et al., 2017). Consistent with the expression profile of *PLC* (Figure 1E), the GFP signals disappeared after far-red irradiation (Figure 1G). These observations indicate that storage vacuole-associated PLC is rapidly silenced following phyA activation.

### *PLC* Negatively Regulates Germination

The expression pattern of *PLC* inspired us to genetically investigate its role in germination. We employed the previously characterized *Arabidopsis* mutant with an intronic T-DNA insertion (Dong et al., 2005) and named this knockdown allele *plc-1* (Supplemental Figure 1A). We also generated a deletion mutant using the CRISPR/Cas9 system with paired guide RNAs. The resulting homozygous mutant, containing a 506 bp deletion that spans the entire coding region, was named *plc-2* (Supplemental Figure 1A). RT-qPCR confirmed that the *plc-2* allele had essentially undetectable *PLC* transcript level in comparison to the wild type (Supplemental Figure 1B). Consistent with previous characterization of *plc-1* (Dong et al., 2005), we found that both *plc* mutants exhibited no apparent difference in the appearance of mature seed compared to the wild type (Supplemental Figure 1C). However, in contrast to the wild type seed that failed to germinate in phyA_OFF_, germination frequency of *plc-1* and *plc-2* significantly increased to 17.3% and 23.3% by 120 h in phyA_OFF_, respectively (Figure 2A and 2B). The approximately 40% germination rate of wild type seed in phyA_ON_ was significantly elevated to 48.0% and 62.7% for *plc-1* and *plc-2*, respectively (Figure 2A and 2C). These results indicate that *PLC* is necessary for effective inhibition of germination in both phyA_OFF_ and phyA_ON_.

**Figure 2.**
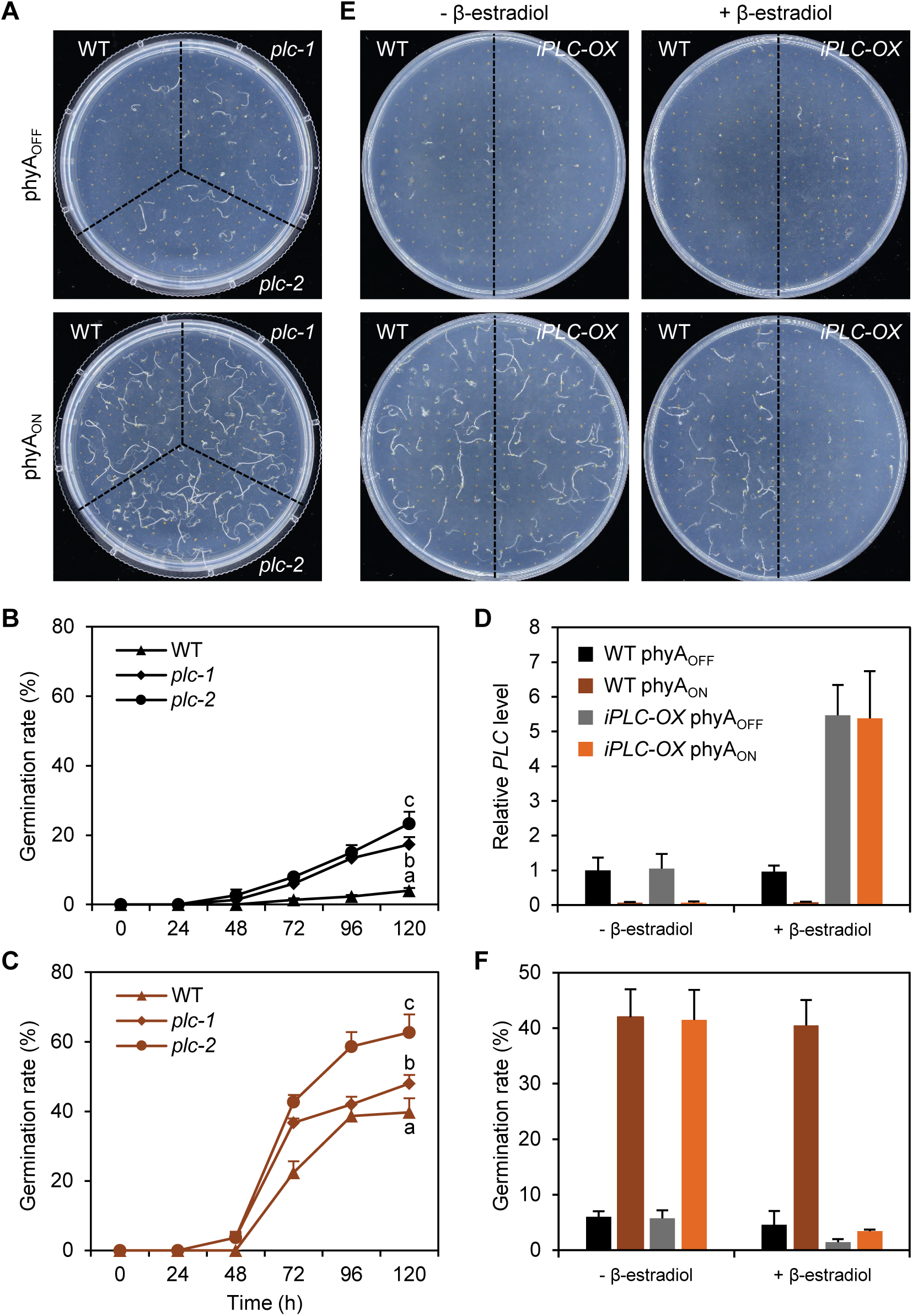
*PLC* Negatively Regulates Germination. (**A**) Representative plates showing germination state of the wild type and *plc* seeds in phyA_OFF_ (top) and phyA_ON_ (bottom). (**B-C**) Quantification of the germination rate over the time course of phyA_OFF_ (B) and phyA_ON_ (C). Data are mean ± SD (n = 3). Different letters represent genotypes with significant differences at 120 h (ANOVA, *p* < 0.05). (**D**) RT-qPCR analysis of relative *PLC* transcript level in the indicated genotypes without and with the application of β-estradiol. Data are mean ± SD (n = 3). (**E**) Representative plates showing germination state of the wild type and *iPLC-OX* seeds in phyA_OFF_ and phyA_ON_ under the indicated treatments. (**F**) Quantification of germination rates of the wild type and *iPLC-OX* seeds. Data are mean ± SD (n = 3).

Moreover, we generated the *iPLC-OX* transgenic *Arabidopsis* plants in which the expression of *PLC* was under the control of a β-estradiol-inducible promoter (Zuo et al., 2000). Mature seed of this line also exhibited no morphological difference from the wild type (Supplemental Figure 1C). The *PLC* transcript was induced to high levels in the *iPLC-OX* seed under both phyA_OFF_ and phyA_ON_ but maintained its normal pattern in the wild type after the application of β-estradiol (Figure 2D; Supplemental Figure 1D). We found that β-estradiol treatment significantly reduced germination rate of the *iPLC-OX* seed, but not of the wild type, in both phyA_OFF_ and phyA_ON_ (Figure 2E and 2F). These results indicate that *PLC* is sufficient for inhibiting germination.

Young seedlings germinated in darkness switch to a different growth mode, termed greening, upon exposure to light (Zhong et al., 2009). Visual inspection revealed that *PLC* represses this process as the greening rate of *plc-2* (referred to as *plc* hereafter) was significantly higher than that of the wild type after dark-germinated seedlings were exposed to white light for 24 h (Figure 3A and 3B). Fluorescence spectral analysis confirmed that the levels of both chlorophylls and the precursor protochlorophyllide in *plc* seedlings were higher than those in the wild type (Figure 3C). Collectively, these results established *PLC* as a negative regulator for germination and the ensued post-germinative growth in light.

**Figure 3.**
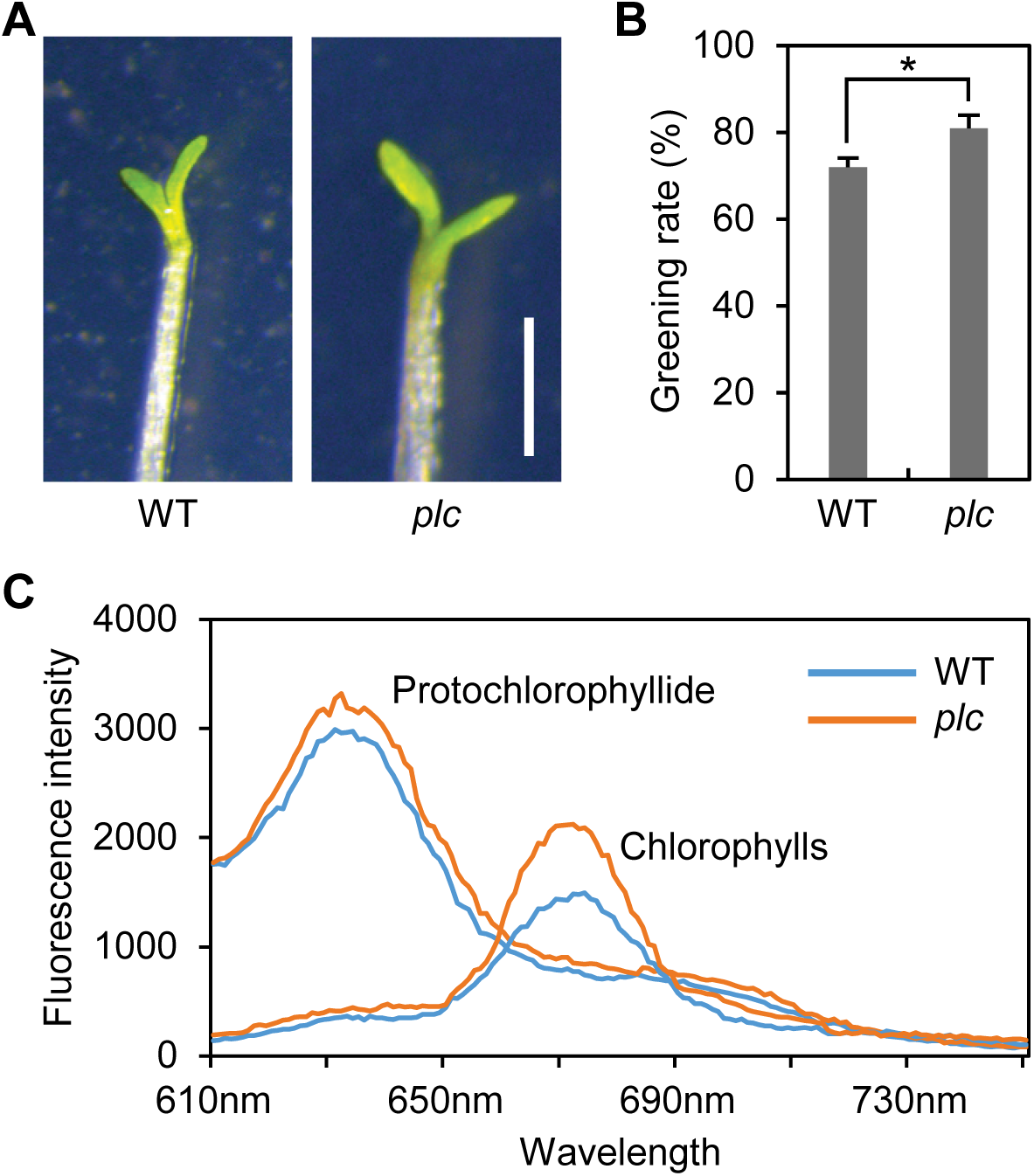
*PLC* Negatively Regulates Seedling Greening. (**A**) Representative wild type and *plc* seedlings that were grown in the dark for 96 h and then exposed to continuous white light for 24 h. Bar, 1 mm. (**B**) Quantified greening rate. Data are mean ± SD (n = 50). *, *p* < 0.05 by Student’s *t* test. (**C**) Comparison of pigment profile in the *plc* and wild type seedlings. Etiolated seedlings grown in the dark for 96 h were assayed for protochlorophyllide by spectral analysis. Chlorophylls were assayed in etiolated seedlings exposed to white light for 24 h.

### *PLC* Is Silenced by miR408 during Germination

In our quest of identifying upstream regulators for *PLC*, we noted that *PLC* is a proven target of miR408 (Abdel-Ghany and Pilon, 2008; Zhang and Li, 2013). Using the standard assay of 5’ RNA ligation-based amplification of cDNA ends, we reassured miR408-guided cleavage of *PLC* mRNA in imbibed seed, which occurred between the 10^th^ and 11^th^ nucleotides in the miR408 recognition site (Figure 4A). From degradome sequencing data for young seedlings, we retrieved reads mapped to the predicted miR408 recognition site in *PLC* (Figure 4B), further confirming miR408 mediated cleavage of the *PLC* transcript. RT-qPCR analysis showed that miR408 level in the seed was stable in the phyA_OFF_ regime but drastically elevated 2 h after the second far-red irradiation and peaked at 8 h in phyA_ON_ (Figure 4C). Thus, miR408 and *PLC* exhibit reciprocal expression pattern following phyA activation (compare Figure 1E and 4C).

**Figure 4.**
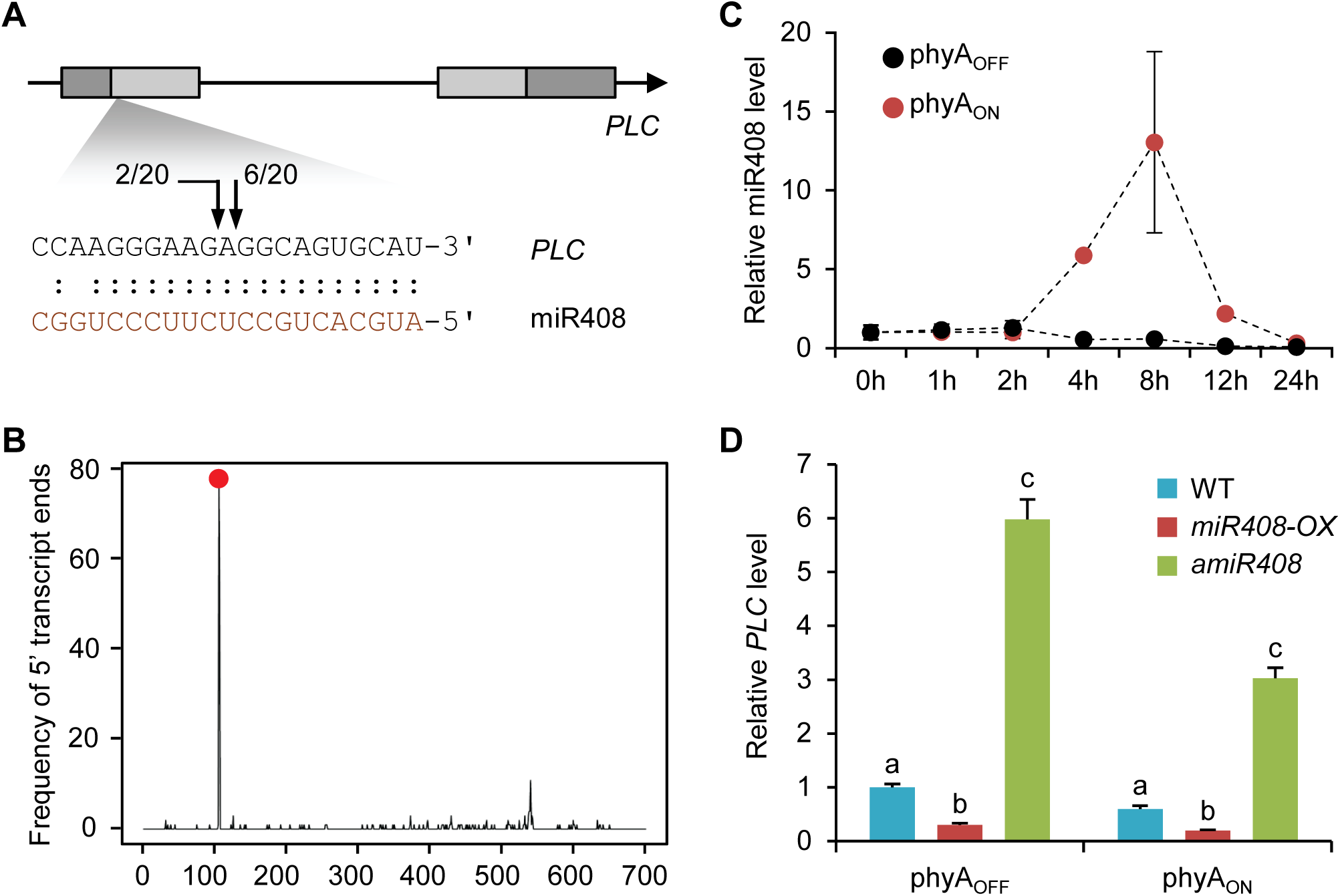
miR408 Represses *PLC* Expression during Germination. (**A**) Confirmation of miR408 targeting on *PLC* in the seed by RNA-ligation based amplification of cDNA ends. Gene structure of *PLC* is shown on top. Base pairing between miR408 and *PLC* is shown on bottom. Arrows mark the detected transcript ends along with frequency of the corresponding clones. (**B**) Degradome sequencing data supporting miR408-guided cleavage of *PLC*. Frequency of the sequenced 5’ ends is plotted against nucleotide position in the *PLC* transcript. Red dot indicates position of the miR408 recognition site. (**C**) RT-qPCR analysis of relative miR408 levels over the time course of phyA_OFF_ and phyA_ON_. Data are mean ± SD (n = 3). (**D**) Relative *PLC* transcript levels in seeds of the indicated genotypes under phyA_OFF_ and phyA_ON_. Data are means ± SD (n = 3). Different letters denote genotypes with significant difference (ANOVA, *p* < 0.05).

To test the effect of miR408 on *PLC* expression during germination, we employed the miR408-overexpressing line (*miR408-OX*) in which the enhanced Cauliflower Mosaic Virus *35S* promoter was used to drive miR408 expression and the miR408-silencing line (*amiR408*) generated by the artificial miRNA method (Zhang and Li, 2013; Zhang et al., 2014). In the *miR408-OX* and *amiR408* seeds, expression of *PLC* in phyA_OFF_ was significantly decreased and increased in comparison to the wild type, respectively (Figure 4D). In phyA_ON_, while *PLC* level was generally lowered compared to that in phyA_OFF_, the trend of relative *PLC* abundance in the wild type, *miR408-OX*, and *amiR408* seeds maintained the same (Figure 4D). Taken together, these results indicate that miR408 negatively modulates *PLC* transcript level through the canonical transcript cleavage mechanism during early seed germination.

### miR408 Is a Positive Regulator of Germination

To investigate the role of miR408 in germination, we examined phenotypes of the *miR408-OX* and *amiR408* seeds (Figure 5A). In phyA_OFF_, the wild type and *amiR408* seeds both failed to germinate while the germination frequency of *miR408-OX* increased to about 80% over the time course (Figure 5B). In phyA_ON_, virtually 100% of the *miR408-OX* seed germinated after 120 h (Figure 5C). In contrast, the germination frequency of *amiR408* only reached approximately 30% after 120 h in phyA_ON_, significantly lower than that of the wild type (Figure 5C). Additionally, we found that miR408 promotes the greening process as the greening rate of *miR408-OX* and *amiR408* was significantly higher and lower than that of the wild type, respectively (Figure 5D to 5F). These results demonstrate miR408 as a positive regulator for germination and post-germinative growth in light.

**Figure 5.**
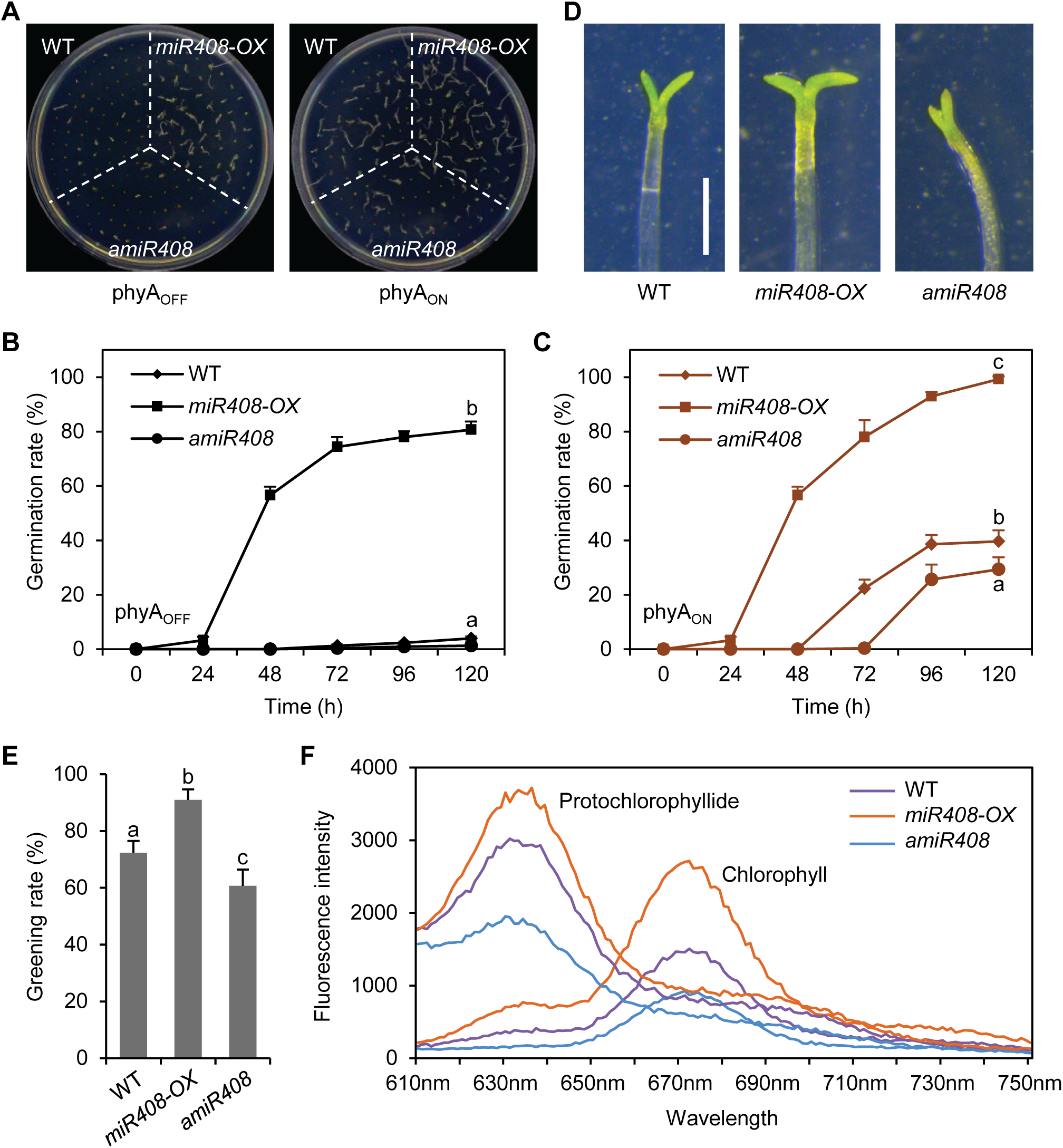
miR408 Promotes Seed Germination and Seedling Greening. (**A**) Representative plates showing germination state of the indicated genotypes under phyA_OFF_ (left) and phyA_ON_ (right). (**B-C**) Quantification of germination rate of the indicated genotypes over the time course of phyA_OFF_ (B) and phyA_ON_ (C). Data are mean ± SD (n = 3). Different letters denote genotypes with significant differences at 120 h (ANOVA, *p* < 0.05). (**D**) Representative etiolated seedlings of the indicated genotypes exposed to white light for 24 h. Bar, 1 mm. (**E**) Quantified greening rate. Data are mean ± SD (n = 50). Different letters represent significant differences (ANOVA, *p* < 0.05). (**F**) Comparison of pigment profile in the indicated genotypes. Protochlorophyllide and chlorophylls were analyzed in etiolated seedlings and etiolated seedlings exposed to white light for 24 h, respectively.

Because the phenotype of *miR408-OX* was stronger than that of *plc*, we examined other miR408 target genes in germination. In *Arabidopsis*, miR408 has four validated targets that all encode cuproproteins, including *PLC, LACCASE 3, 12*, and *13* (Abdel-Ghany and Pilon, 2008; Zhang and Li, 2013; Zhang et al., 2014). Based on known expression profiles (Winter et al., 2007; Zhuang et al., 2020), the three *LAC* genes did not exhibit substantial expression in the seed (Supplemental Figure 2A). Phenotypic comparison of the *lac3, lac12, lac13*, and *plc* single mutants revealed that only *plc* displayed significantly elevated germination frequency than the wild type in both phyA_OFF_ and phyA_ON_ (Supplemental Figure 2B). We further generated the *lac12 lac13* double mutant and the *plc lac12 lac13* triple mutant. We found that the *plc lac12 lac13* seed exhibited the same germination phenotype as *plc* while *lac12 lac13* showed no difference from the wild type (Supplemental Figure 2C). These results indicate that other miR408 target genes are not involved in germination. Thus, whether the relatively weaker phenotype of *PLC* loss-of-function was due to the compensation by other phytocyanins warrants further investigation.

### PIF1 Directly Suppresses *MIR408* to Promote *PLC* Expression

Our next goal was to find out how the miR408-*PLC* module is regulated by light signaling. Previously, we reported that ELONGATED HYPOCOTYL 5 (HY5), by binding to the G-box in the *MIR408* promoter, activates *MIR408* in response to increasing light irradiation (Zhang et al., 2014). Since PIFs and HY5 could antagonistically adjust the expression of common target genes (Chen et al., 2013; Toledo-Ortiz et al., 2014; Shi et al., 2018), we investigated the possibility that PIF1 transcriptionally represses *MIR408*. Using PIF1 chromatin immunoprecipitation (ChIP) sequencing data (Pfeiffer et al., 2014), we identified a significant PIF1 binding peak in the *MIR408* promoter encompassing the G-box (Figure 6A). Using the *PIF1-OX* line expressing MYC-tagged PIF1 driven by the *35S* promoter (Oh et al., 2004), we performed ChIP with an anti-MYC antibody. qPCR analysis revealed that PIF1 occupancy at the G-box containing DNA fragment was enriched over fivefold in *PIF1-OX* relative to the wild type (Figure 6B), confirming *PIF1* as a direct upstream regulator of *MIR408*.

**Figure 6.**
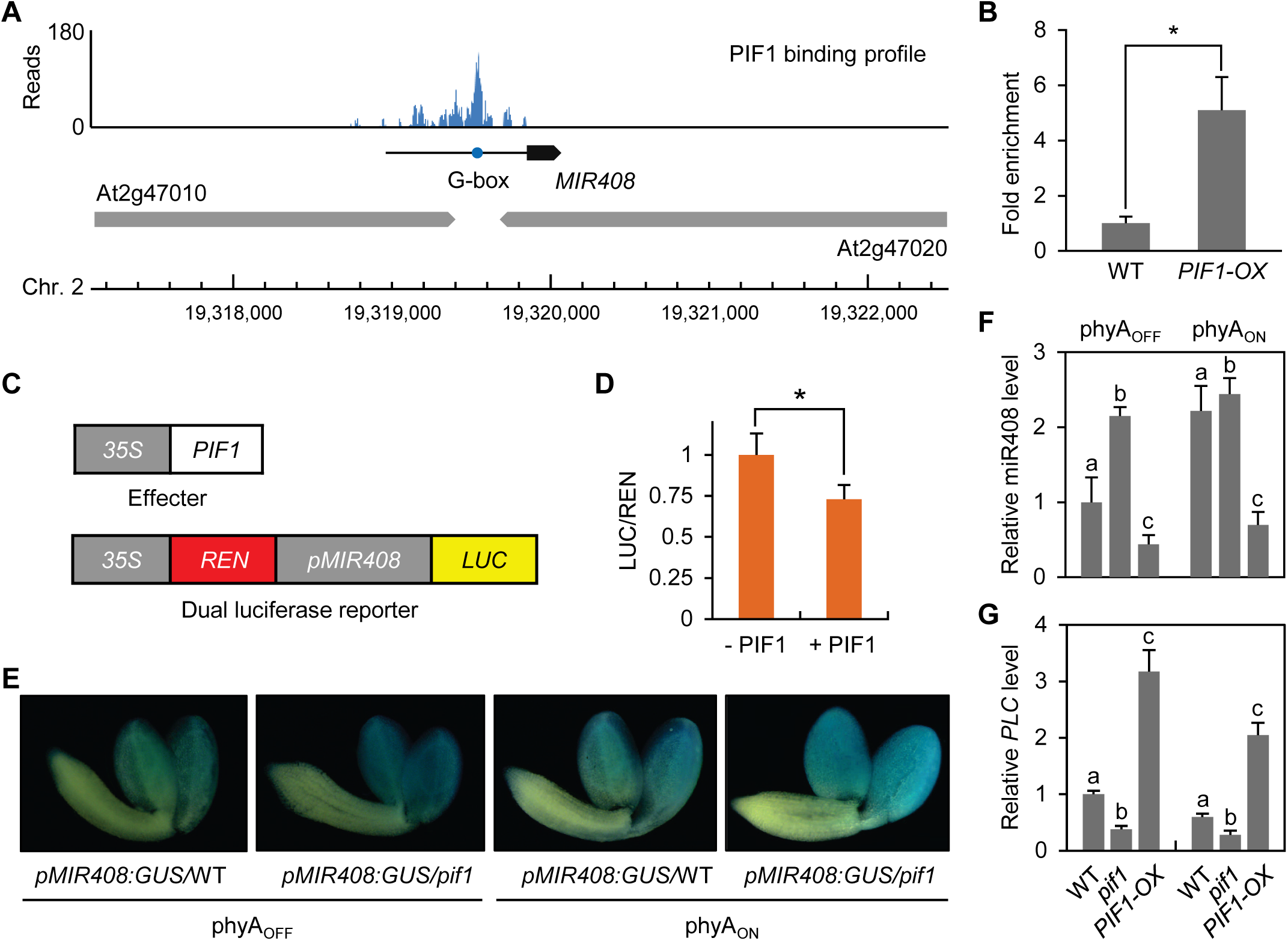
PIF1 Suppresses miR408 Expression by Binding to the *MIR408* Promoter. (**A**) PIF1 occupancy profile at the *MIR408* locus. Significantly enriched PIF1 ChIP-sequencing reads were obtained from Pfeiffer et al. (2014) and mapped onto the *Arabidopsis* genome coordinates. Loci are represented by block arrows. The blue circle marks the G-box (CACGTG) in the *MIR408* promoter (horizontal line). (**B**) ChIP-qPCR confirming PIF1 binding to the *MIR408* promoter. An anti-MYC antibody was used to precipitate chromatin from *PIF1-OX* and wild type seeds. Enrichment of PIF1 binding was determined by qPCR analysis. Data are mean ± SD (n = 3). *, *p* < 0.05 by Student’s *t* test. (**C**) Transient dual luciferase assay showing PIF1 repression of *MIR408*. The *pMIR408:LUC* reporter concatenated to *35S:REN* was used to transform tobacco protoplasts with either the empty vector (-PIF1) or a PIF1-expressing construct (+ PIF1). (**D**) Quantification of the LUC/REN luminescence ratio. Data are mean ± SD (n = 3). *, *p* < 0.05 by Student’s *t* test. (**E**) Comparison of GUS activity in transgenic seed expressing *pMIR408:GUS* in the wild type or *pif1* background in phyA_OFF_ and phyA_ON_. Bar, 500 μm. (**F-G**) RT-qPCR analysis of relative miR408 (F) and *PLC* (G) transcript levels. Data are mean ± SD (n = 3). Different letters denote genotypes with significant differences (ANOVA, *p* < 0.05).

To ascertain the net effect of PIF1 on the *MIR408* promoter, we employed the firefly luciferase (LUC) and *Renilla* luciferase (REN) dual reporter system (Hellens et al., 2005). We generated the *pMIR408:LUC* reporter and the *35S:PIF1* effector constructs (Figure 6C).

Following transfection of tobacco leaf protoplasts, we found that co-expression of *PIF1* significantly reduced the LUC/REN ratio (Figure 6D), indicating that PIF1 negatively modulates the *MIR408* promoter. To corroborate this relationship in *Arabidopsis*, we fused the *β-glucuronidase* (*GUS*) coding region with the *MIR408* promoter and expressed the same reporter gene in either the wild type (*pMIR408:GUS/*WT) or *pif1* (*pMIR408:GUS*/*pif1*) background (Figure 6E). We found that GUS activity, mainly detected in the cotyledons of imbibed seed, was higher in the *pif1* background in both phyA_OFF_ and phyA_ON_ (Figure 6E), confirming *PIF1*-mediated suppression of *MIR408 in planta*.

To monitor the influence of *PIF1* on miR408 and *PLC* transcript accumulation, we performed RT-qPCR analysis of the *pif1* and *PIF1-OX* lines. This analysis revealed that miR408 abundance significantly increased in *pif1* but decreased in *PIF1-OX* in both phyA_OFF_ and phyA_ON_ with reference to the wild type (Figure 6F). Conversely, *PLC* abundance significantly increased in *PIF1-OX* but decreased in *pif1* in both phyA_OFF_ and phyA_ON_ (Figure 6G). Taken together, these results demonstrate that PIF1 binds to the *MIR408* promoter and represses accumulation of miR408 in the seed, which in turn post-transcriptionally silences *PLC*, thereby forming a *PIF1-MIR408-PLC* repression cascade.

### *PIF1, MIR408*, and *PLC* Act in the Same Pathway to Regulate Germination

Consistent with previously reports (Oh et al., 2004), we found that *pif1* seed completely germinated by 72 h in both phyA_OFF_ and phyA_ON_ (Figure 7A). This enhanced germination phenotype was observed for the *miR408-OX* and *plc* seeds as well (Figure 2B and 5B).

**Figure 7.**
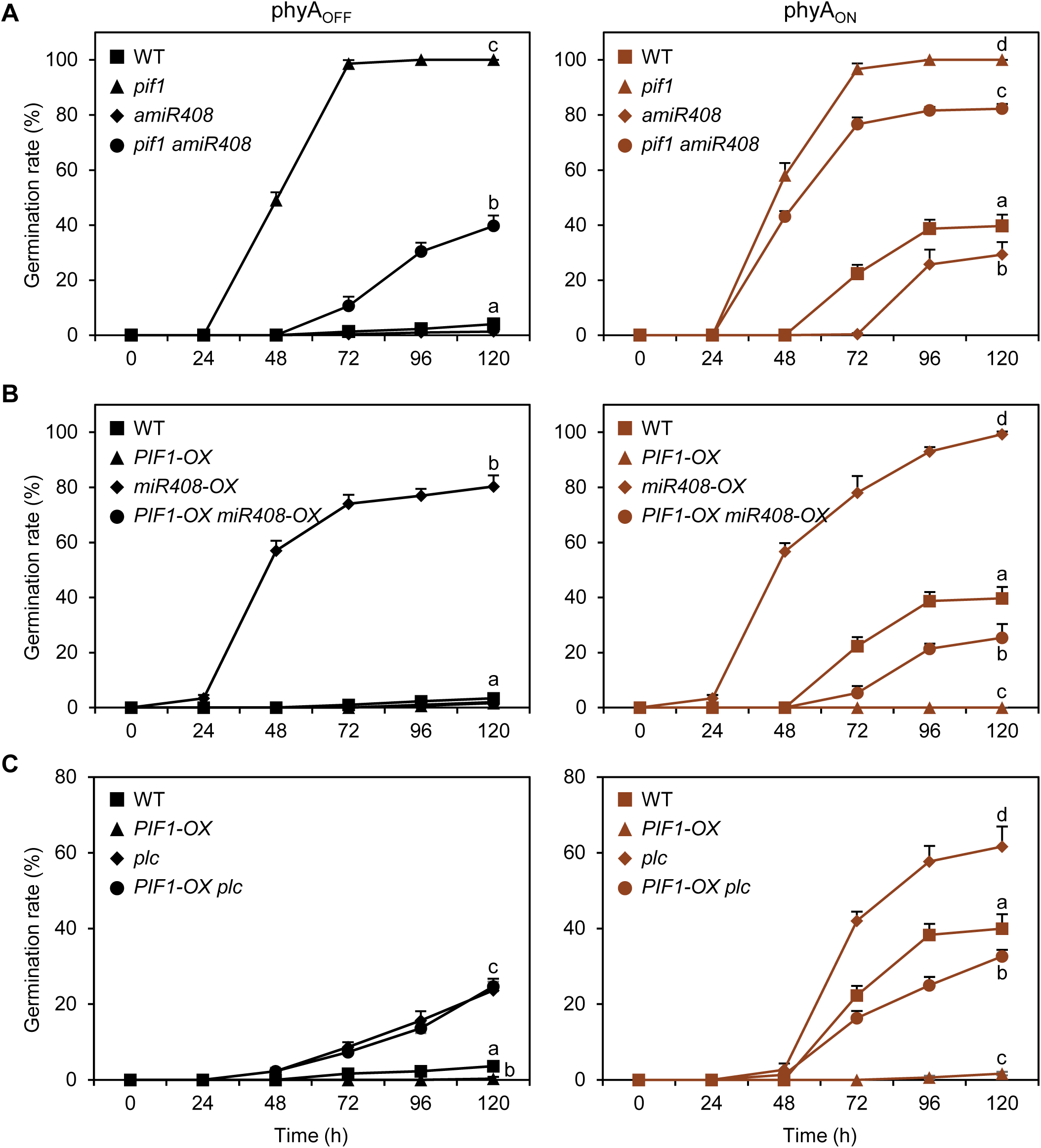
Genetic Analysis of the *PIF1-MIR408-PLC* Pathway. (**A**) The *amiR408* line was crossed with *pif1* to generate the *pif1 amiR408* double mutant. Seeds from these lines and the wild type were assayed for germination rates over the time course of phyA_OFF_ (left) and phyA_ON_ (right). (**B**) The *miR408-OX* line was crossed with *PIF1-OX* to generate the *PIF1-OX miR408-OX* double over-expression line. Seeds from these lines and the wild type were assayed for germination rates in phyA_OFF_ and phyA_ON_. (**C**) Comparison of germination rates of the *PIF1-OX, plc*, and *PIF1-OX plc* seeds. Data are all means ± SD (n = 3). Different letters denote genotypes with significant differences at 120 h (ANOVA, *p* < 0.05).

Conversely, *PIF1-OX* seed failed to germinate even by 120 h in phyA_ON_ (Figure 7B). We found that *amiR408* and β-estradiol treated *iPLC-OX* seeds exhibited similar phenotypes like *PIF1-OX* (Figure 2F and 5C). These results indicate that the molecularly delineated *PIF1-MIR408-PLC* repression cascade was in line with the germination phenotypes of the relevant mutants.

To further confirm that *PIF1, MIR408*, and *PLC* acts in the same genetic pathway, we generated and analyzed three double mutants involving *pif1* and *PIF1-OX*. We found that germination rate of the *pif1 amiR408* seed was substantially reduced compared to that of *pif1* (Figure 7A). By 120 h in phyA_OFF_ and phyA_ON_, the near complete germination of *pif1* was lowered to about 40% and 80% by *amiR408*, respectively (Figure 7A). Regarding the *PIF1-OX miR408-OX* double overexpression seed, over 20% germinated by 120 h in phyA_ON_ (Figure 7B), indicating that *miR408-OX* was able to partially rescue the germination defect of *PIF1-OX*. These results demonstrate that *MIR408* is downstream of *PIF1* in the same pathway. We also tested the genetic relationship between *PLC* and *PIF1* by generating the *PIF1-OX plc* line. We found that the germination profile of this line was similar to that of the wild type with substantially increased rates than *PIF1-OX* in both phyA_OFF_ and phyA_ON_ (Figure 7C), indicating that *PLC* is also downstream of *PIF1*. Thus, *PIF1, MIR408*, and *PLC* act sequentially in the same pathway to regulate seed germination.

### *PIF1, MIR408*, and *PLC* Regulate Overlapping Cohorts of Genes

As a master transcriptional regulator, PIF1 programs the seed germination related transcriptome (Oh et al., 2009; Shi et al., 2013; Pfeiffer et al., 2014). To test whether that transcriptome is regulated through the *PIF1-MIR408-PLC* pathway, we performed RNA-sequencing analysis of the wild type, *pif1*, and *miR408-OX* seeds imbibed in darkness. The *iPLC-OX* imbibed seed with or without β-estradiol treatment was also analyzed. Consistent with previous reports (Oh et al., 2009; Shi et al., 2013), we identified 5,640 genes that were differentially expressed between *pif1* and the wild type, which was defined as the *PIF1*-regulated set (Figure 8A). Genes differentially expressed between *miR408-OX* and the wild type were defined as the *MIR408*-regulated set, which included 4,294 genes (Figure 8A). Differentially expressed genes in *iPLC-OX* with and without β-estradiol treatment were defined as the *PLC*-regulated set, which included 14,646 genes (Figure 8A).

**Figure 8.**
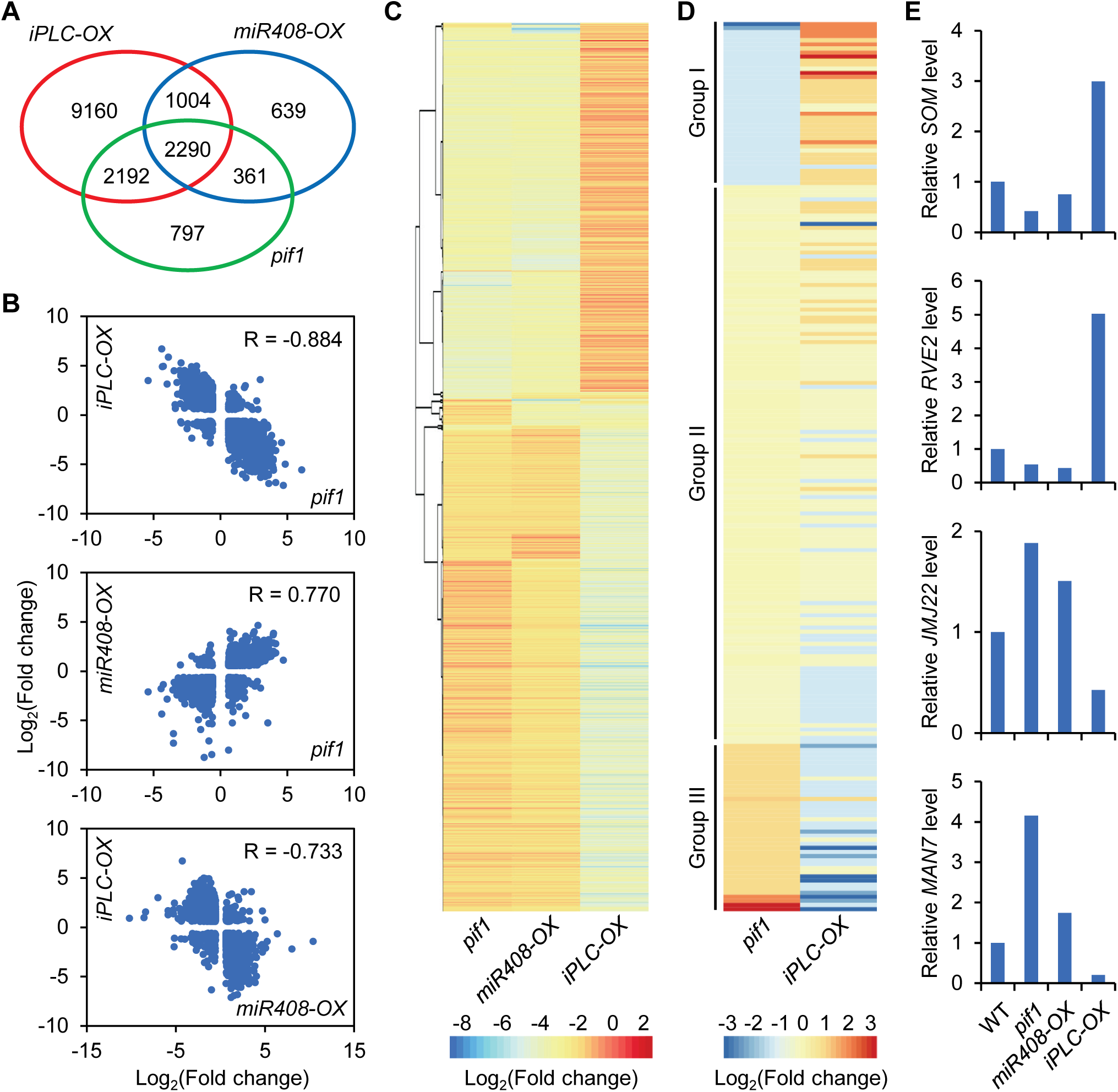
Transcriptomic Analysis of the *PIF1-MIR408-PLC* Pathway. (**A**) Venn diagram showing the relationships of *PIF1, MIR408*, and *PLC* regulated genes. Differentially expressed genes were identified from RNA-sequencing analysis of *pif1, miR408-OX*, and *iPLC-OX* seeds against the respective controls. (**B**) Scatterplots showing pairwise correlation of the relative expression levels of the three sets of coregulated genes in *pif1, miR408-OX*, and *iPLC-OX* against the respective controls. R, Pearson correlation coefficient. (**C**) Hierarchical clustering of the 2,290 genes differentially expressed in *pif1, miR408-OX*, and *iPLC-OX* against the respective controls. Colors represent the Log_2_ transformed fold change. (**D**) Clustering analysis of the 218 genes associated with the GO term “seed germination” (GO:0009845). The genes were divided in three grouped based relative expression level in *pif1* against the wild type. Group I, repressed in *pif1*; Group II, not differentially expressed; Group III, induced in *pif1*. (**E**) Expression pattern of representative Group I (*SOM* and *RVE2*) and Group III (*JMJ22* and *MAN7*) genes in the indicted RNA-sequencing datasets.

Venn diagram analysis revealed that there were 2,651 common genes between the *PIF1*-regulated and the *MIR408*-regulated sets, 4,482 common between the *PIF1*-regulated and the *PLC*-regulated sets, and 3,294 common between the *MIR408*-regulated and the *PLC*-regulated sets (Figure 8A). Based on Pearson correlation coefficient of fold changes against the controls, we found that these common genes exhibited high pairwise correlations between the compared genotypes (Figure 8B). Venn diagram analysis further revealed 2,290 genes that were differentially expressed in *pif1, miR408-OX*, and *iPLC-OX* compared to the respective controls (Figure 8A). Clustering analysis showed that the vast majority of these genes was regulated in the same direction in *pif1* and *miR408-OX* but the opposite direction in *iPLC-OX* (Figure 8C). Together these results indicate that the *PIF1-MIR408-PLC* pathway regulates large cohorts of common target genes in the seed.

Since 79.5% of the *PIF1*-related genes, or 4,482 out of 5,640, was differentially regulated in *iPLC-OX* (Figure 8A), which also exhibited the highest pairwise correlation among the genotypes (R = 0.884; Figure 8B), we selected the *PIF1-PLC* coregulated genes for further analyses (Supplemental Dataset 1). Gene Ontology (GO) analysis revealed that the *PIF1-PLC* coregulated genes were preferentially associated with terms in two categories: seed development and germination, and hormone metabolism and signaling (Supplemental Figure 3). For example, the GO term “seed germination” was associated with 218 genes in *Arabidopsis* (Supplemental Dataset 2). These genes could be divided into three groups based on their expression pattern in *pif1* (Figure 8D). Genes in group I (40 out of 218, or 18.3%) and III (41 out of 218, or 18.8%) were substantially down-regulated and up-regulated in *pif1*, respectively. They were reversely modulated in *iPLC-OX* (Figure 8D).

Further inspection revealed that many individual genes that have been genetically or functionally implicated in germination related processes were included in group I and III (Supplemental Figure 4). For example, *SOM* and *RVE2* from group I were reported to inhibit light dependent germination (Kim et al., 2008; Jiang et al., 2016). On the contrary, *JMJ22* and *MAN7* from group III were reported to promote seed germination (Iglesias-Fernández et al., 2011; Cho et al., 2012). These two types of genes were regulated in an opposite pattern in *pif1* and *iPLC-OX* (Figure 8E). Collectively, these results indicate that *PLC* is a key node downstream of *PIF1* to mediate the transcriptomic changes underlying light dependent germination.

### The *PIF1-MIR408-PLC* Pathway Regulates GA and ABA Biosynthesis

As rapid removal of PIF1 is critical for ultimately establishing the high-GA-low-ABA state, we examined whether PLC impacts the GA-ABA balance. To this end, we first inspected gene expression profile along the GA metabolic pathway. This analysis revealed that the *PIF1-MIR408-PLC* pathway regulates in particular *GA3ox1* and *GA2ox2*, which encode the GA3-oxidase that catalyzes the terminal GA biosynthetic step and the GA2-oxidase that catabolizes bioactive GAs, respectively (Figure 9A) (Yamaguchi, 2008). This finding was corroborated by RT-qPCR analysis showing that *GA3ox1* was up-regulated in *pif1, miR408-OX*, and *plc*, but down-regulated in *PIF1-OX* and *amiR408* seeds compared to the wild type in both phyA_OFF_ and phyA_ON_ (Figure 9B). In contrast, *GA2ox2* was up-regulated in *PIF1-OX* and *amiR408* but down-regulated in *pif1, miR408-OX*, and *plc* seeds relative to the wild type (Figure 9B). These results indicate that the *PIF1-MIR408-PLC* pathway targets the *GA3ox* and *GA2ox* steps in GA biosynthesis to regulate the level of bioactive GA.

**Figure 9.**
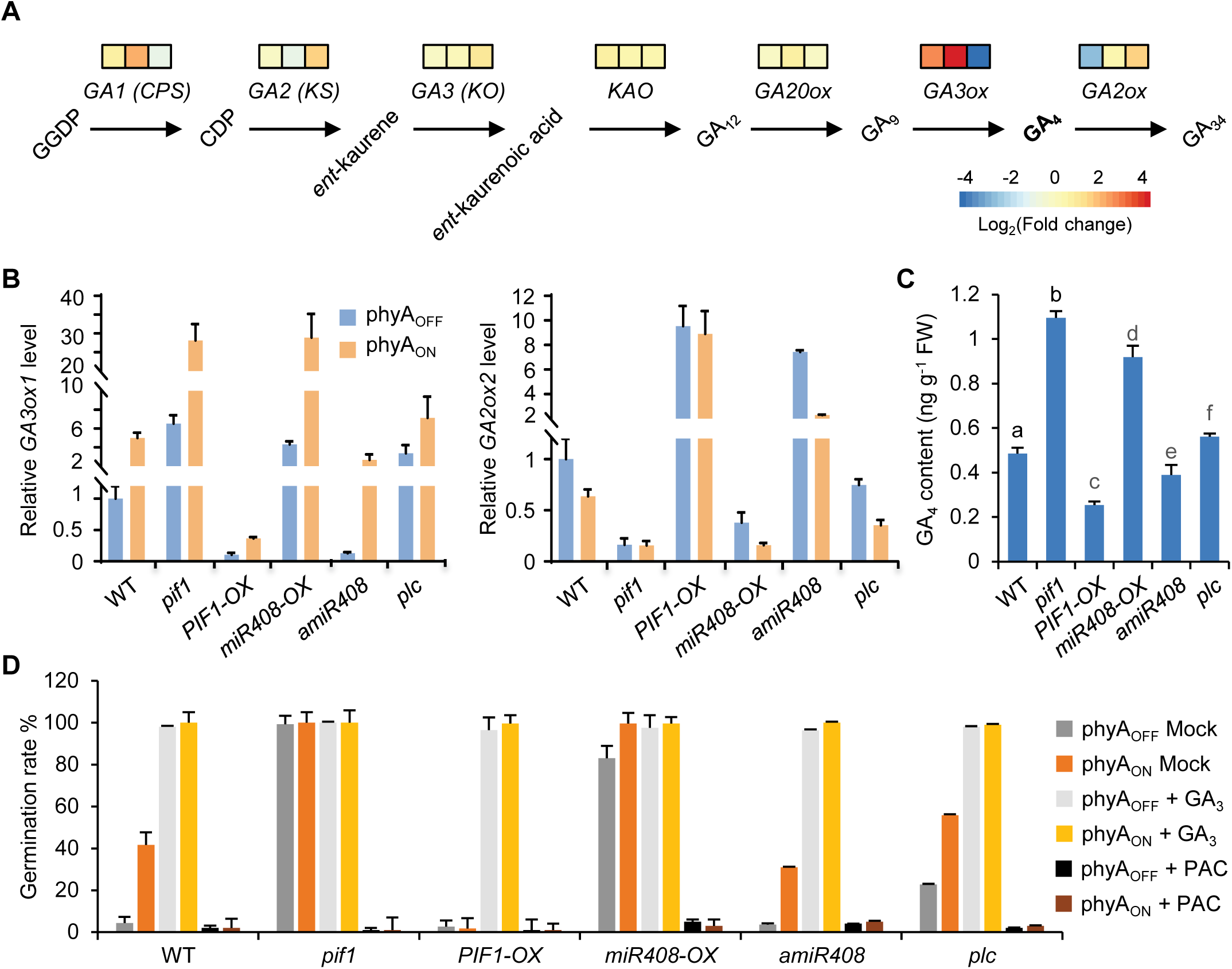
The *PIF1-MIR408-PLC* Pathway Modulates GA Biosynthesis. (**A**) Diagram of a simplified GA biosynthesis pathway illustrating genes influenced by the *PIF1-MIR408-PLC* pathway. Genes associated with the individual biosynthesis steps are shown on top of the arrows. Colored boxes indicate relative expression levels of the corresponding gene in *pif1, miR408-OX*, and *iPLC-OX* against the respective controls. (**B**) RT-qPCR analysis of relative transcript level of the *GA3ox1* and *GA2ox2* genes in the indicated seeds under phyA_OFF_ and phyA_ON_. Data are mean ± SD (n = 3). (**C**) Quantification of endogenous GA_4_ levels in imbibed seed of the indicated genotypes. Data are mean ± SD (n = 3). Different letters denote genotypes with significant differences (ANOVA, *p* < 0.05). (**D**) Germination rates of the indicated seeds in phyA_OFF_ and phyA_ON_ with different treatments. Mock, no chemical treatment; GA_3_, 10 μM GA_3_; PAC, 100 μM paclobutrazol. Data are mean ± SD (n = 3).

Next, we directly quantified the amount of bioactive GA in the seed using the ultrahigh-performance liquid chromatography-triple quadrupole mass spectrometry (UPLC-MS/MS) method (Fu et al. 2012; Ma et al., 2015). Compared to the wild type, level of GA_4_, the major bioactive GA in *Arabidopsis* seed (Oh et al., 2006), was significantly elevated in *pif1, miR408-OX*, and *plc* seeds (Figure 9C). In *PIF1-OX* and *amiR408*, GA_4_ level was significantly reduced compared to the wild type (Figure 9C). These results indicate that the *PIF1-MIR408-PLC* pathway is capable of modulating the level of bioactive GA in the seed.

We further performed pharmacological analyses using bioactive GA_3_ and paclobutrazol, an inhibitor of GA biosynthesis. We found that GA_3_ application promoted all seeds, including *PIF1-OX* and *amiR408*, to complete germination in phyA_OFF_ (Figure 9D). Conversely, paclobutrazol blocked germination of all seeds, including *pif1, miR408-OX*, and *plc*, in phyA_ON_ (Figure 9D). Taken together, the gene expression, hormone quantification, and pharmacological results demonstrate that the *PIF1-MIR408-PLC* cascade regulates germination by modulating the level of bioactive GA in the seed.

Transcriptomic profiling also revealed that *ABA1, NCED6* and *NCED9* were among the most substantially influenced ABA biosynthetic genes, which were downregulated in *pif1* and *miR408-OX* but upregulated in β-estradiol treated *iPLC-OX* seeds compared to the controls (Figure 10A). RT-qPCR analysis confirmed that *ABA1, NCED6* and *NCED9* were positively regulated by *PIF1* and *PLC* but negatively regulated by miR408 in both phyA_OFF_ and phyA_ON_ (Figure 10B). Chemical quantification showed that endogenous ABA level in *pif1, miR408-OX*, and *plc* seeds was significantly reduced compared to that in the wild type (Figure 10C).

**Figure 10.**
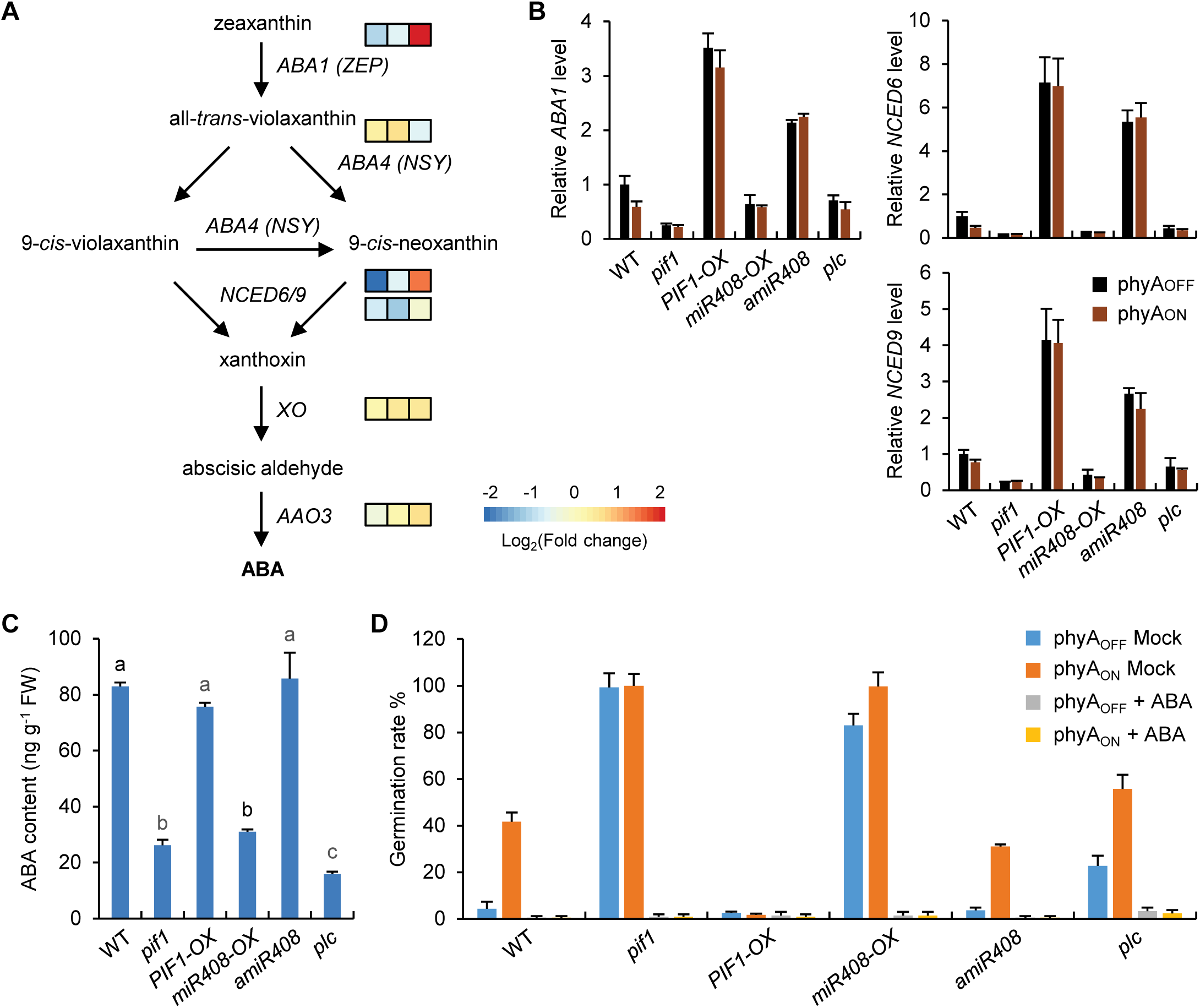
The *PIF1-MIR408-PLC* Pathway Regulates ABA Biosynthesis. (**A**) Diagram of a simplified ABA biosynthesis pathway illustrating genes influenced by the *PIF1-MIR408-PLC* pathway. Colored boxes indicate relative expression levels of the corresponding gene in *pif1, miR408-OX*, and *iPLC-OX* against the respective controls. (**B**) RT-qPCR analysis of the relative transcript levels of ABA metabolic genes *ABA1, NCED6*, and *NCED9* in the indicated seeds. Data are mean ± SD (n = 3). (**C**) Quantification of endogenous ABA level in the indicated seeds. Data are mean ± SD (n = 3). Different letters denote groups with significant differences (ANOVA, *p* < 0.05). (**D**) Germination rate of the indicated seeds in phyA_OFF_ and phyA_ON_ with or without the application of 5 μM ABA.

Furthermore, we found that pharmacological treatment with ABA blocked germination of all seeds, including *pif1, miR408-OX*, and *plc* in phyA_ON_ (Figure 10D). Together, these results indicate that the *PIF1-MIR408-PLC* pathway regulates germination through reciprocally modulating the biosynthesis of GA and ABA.

### PLC Is Conserved in Seed Plants

Finally, to provide phylogenetic evidence supporting PLC as a key node in germination, we examined PLC conservation in seed plants. Searching for putative PLC orthologs from sequenced land plant genomes identified 276 PLC sequences from 52 seed plants but not non-seed plants (Figure 11; Supplemental Figure 5). Comparison of PLC sequences and domain organization to the most homologous blue copper proteins in three representative non-seed plants revealed two salient features of PLC. First, all PLCs were found to contain only a signal peptide at the N-terminus and a type-I copper binding motif at the C-terminus (Figure 11; Supplemental Figure 5). These two domains exhibited an extremely compact organization. For example, the *Arabidopsis* PLC possessed 129 amino acid residues of which 33 (25.6%) were devoted to the signal peptide and 95 (73.6%) to the copper binding motif (Figure 11). Second, a miR408 recognition site was found near the 5’ end of the coding region of all examined *PLC* transcripts (Figure 11), which has been experimentally validated in several plant species (Abdel-Ghany and Pilon, 2008; Zhou et al., 2010; Feng et al., 2013; Zhang and Li, 2013). Moreover, length and domain organization suggest that PLC in *Ginkgo* may resemble the prototype of this protein, which was further evolved in angiosperm by trimming the C-terminus to a bare-bones copper binding motif (Figure 11). Taken together, our results indicate that PLC has specifically evolved in seed plants as a miR408 targeted, storage vacuole associated compact cuproprotein, which balances GA and ABA levels for controlling germination (Figure 12).

**Figure 11.**
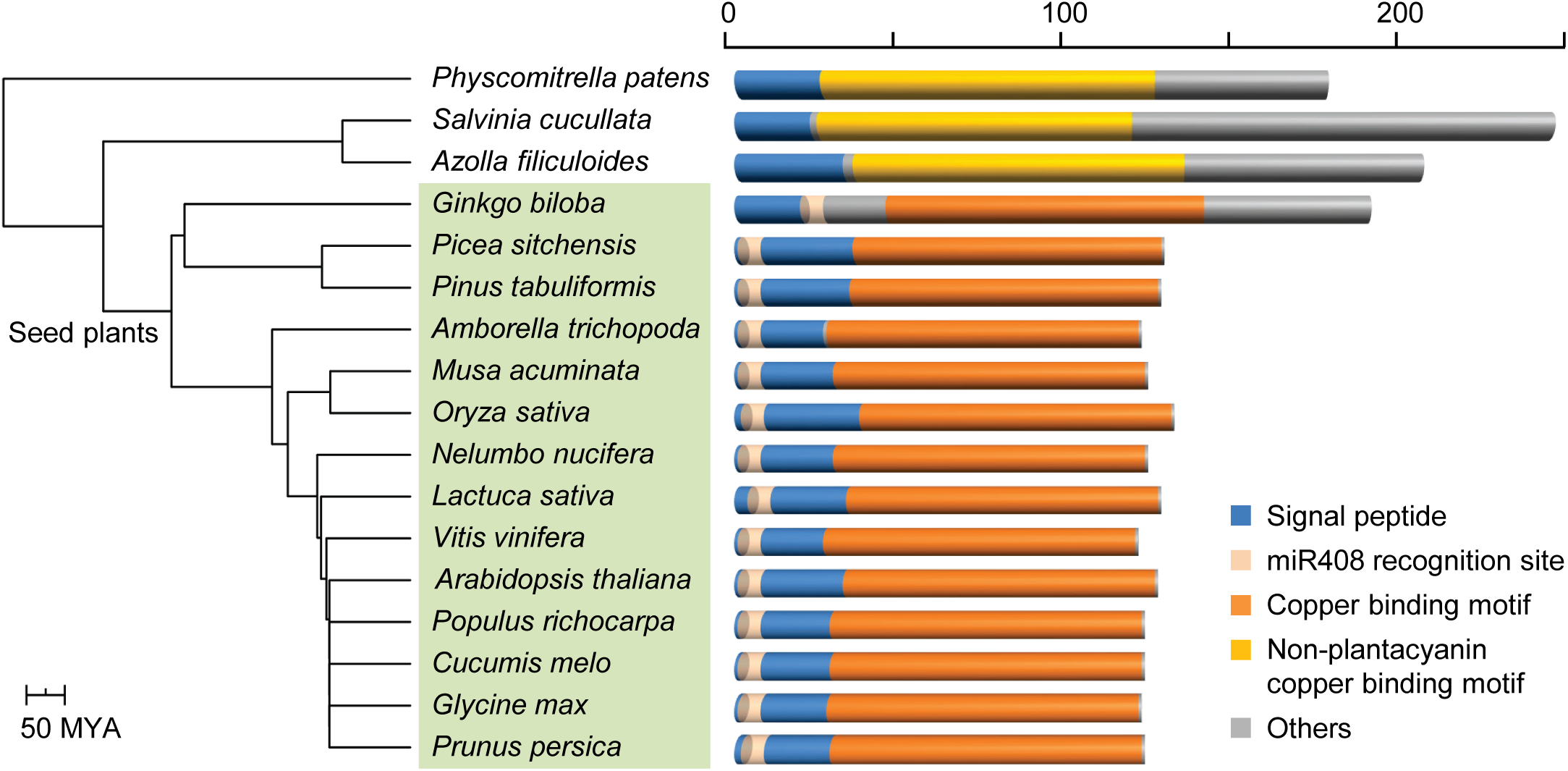
PLC Exhibits Features Conserved in Seed Plants. Comparison of PLC and related blue copper proteins in representative land plants. Shown on the left is a species tree. Branch length reflects evolutionary divergence time in millions of years inferred from TimeTree. Species with identified PLCs are shaded in green. Domains are shown with different colors on the right. Scale represents accumulative number of amino acids.

**Figure 12.**
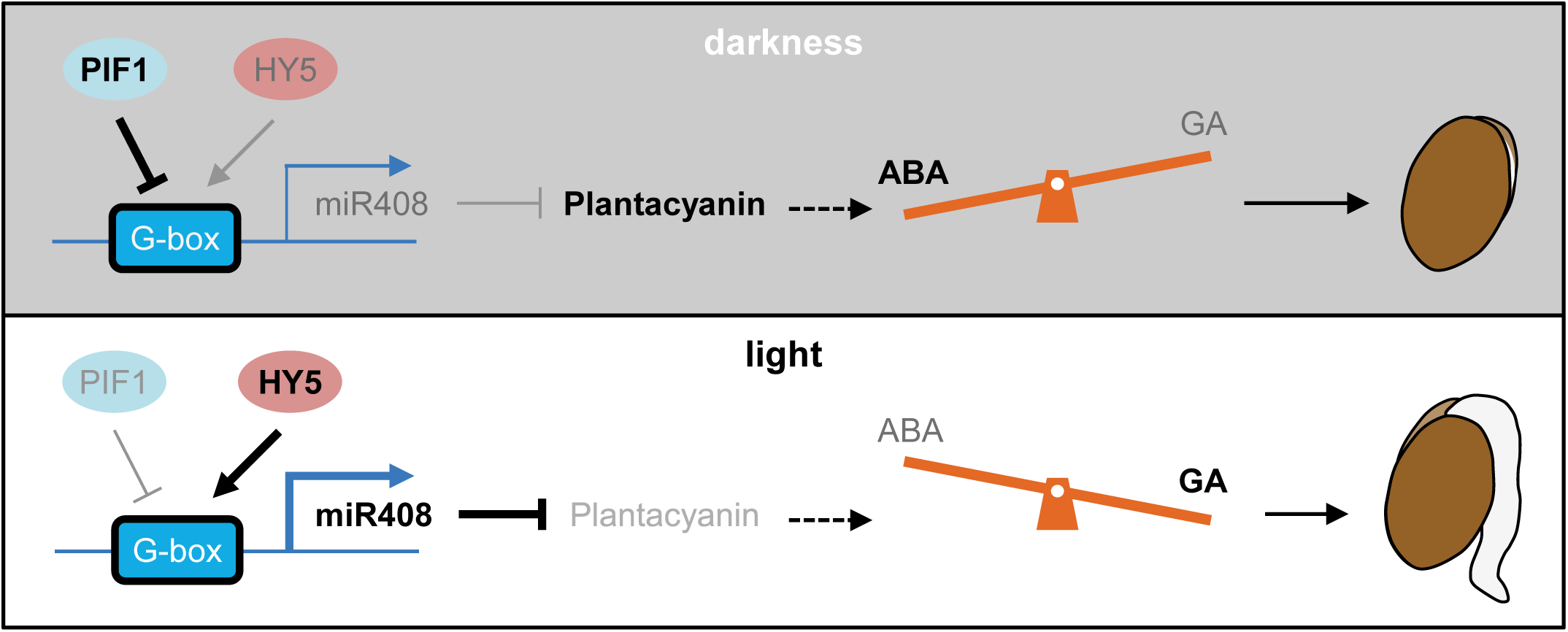
Working Model of Light-dependent Seed Germination Mediated by PLC. The PIF1/HY5-miR408 module is critical for regulating PLC abundance in light-induced seed germination. Both PIF1 and HY5 can bind to the G-box cis-element in the *MIR408* promoter and thereby modulate cellular miR408 level. In darkness, stabilized PIF1 is the predominant regulator leading to transcriptional repression of miR408, which allows PLC to accumulate. Upon light irradiation, HY5 activation and PIF degradation leads to transcriptional de-repression of miR408, which in turn silences *PLC*. Removal of storage vacuole located PLC facilitates establishing the high-GA-low-ABA hormonal profile that eventually sets germination in motion and promotes post-germinative growth in light.

## DISCUSSION

The seeds are equipped with elaborate molecular mechanisms to monitor and transduce the light signals for proper germination, which is vital for survival of seed plants (Oh et al., 2004; Finch-Savage and Leubner-Metzger, 2006; Seo et al., 2009; Shi et al., 2015). Decades of research has shown that a decisive event downstream of light signaling is establishment of the high-GA-low-ABA hormonal state (Nambara and Marion-Poll, 2005; Oh et al., 2006; 2007; Yamaguchi, 2008; Seo et al., 2009; Shu et al., 2016). Elucidating how the light signals are converted into the hormonal profiles is critical to our understanding of seed biology, which bears immediate relevance to agriculture and human nutrition.

### A Long Repression Cascade Regulating Light Dependent Germination

In this study, we used far-red light triggered, phyA dependent germination as the experiment model (Oh et al., 2004; 2006; Cho et al., 2012). Our comprehensive molecular and genetic analyses (Figure 1 to 7), incorporated to established light signaling framework (Castillon et al., 2007; Leivar and Quail, 2010), delineated the phyA-PIF1-miR408-PLC repression cascade as a key regulatory mechanism in germination (Figure 12). Through transcriptome profiling and quantification of endogenous GA and ABA levels, we found that this signal relay chain regulates the conversion of light signals into hormonal profiles (Figure 8 and 10). In darkness, absence of active phyA leads to PIF1 accumulation (Shen et al., 2008), which suppresses transcription of *MIR408* (Figure 6). Low level of miR408 in turn allows PLC to accumulate in the storage vacuole (Figure 1 and 4), which correlates with the low-GA-high-ABA state (Figure 9 and 10). Upon far-red irradiation, phyA is activated to rapidly destabilize PIF1 (Oh et al., 2007; Shen et al., 2008), releasing *MIR408* from transcriptional inhibition (Figure 6). Accumulation of miR408 then leads to *PLC* silencing (Figure 1 and 4), which correlates with the high-GA-low-ABA state (Figure 9 and 10). This chain of events forms a multistep repressor cascade typical for developmental transcription networks, which generates robust temporal delay (Rosenfeld and Alon, 2003; Shoval and Alon, 2010). The phyA-PIF1-miR408-PLC repression cascade therefore may help to specify the time from light perception to PLC turnover.

Based on phylogenetic findings, the phyA-PIF1-miR408-PLC cascade appears to have formed by sequentially adding downstream components during evolution of the seed plants. The phytochrome signaling pathway and the PIF family were shown to originate in the ancestors of charophytes (Han et al., 2019). Inhibition of PIFs by phytochromes to regulate light responses was conserved at least in vascular plants and the liverworts (Lee and Choi, 2017; Han et al., 2019). On the other hand, miR408 is deeply conserved in land plants including moss (Pan et al., 2018; Guo et al., 2020) while PLC acquired the miR408 recognition site after the emergence of seed plants (Figure 11). These observations suggest that the repression cascade has specifically evolved in seed plants, taking advantage of extant regulatory modules, for controlling seed germination.

HY5 and PIFs are known to function antagonistically to adjust the expression of common target genes related to biological processes such as seedling establishment, photosynthetic pigment synthesis, production of reactive oxygen species, and phosphate starvation response (Chen et al., 2013; Toledo-Ortiz et al., 2014; Sakuraba et al., 2018; Shi et al., 2018). We reported previously that HY5 binds to the G-box in the *MIR408* promoter and promotes miR408 accumulation in young seedlings in a light intensity dependent manner (Zhang et al., 2014). The discovery of PIF1 represses *MIR408* transcription via binding to the G-box (Figure 6) indicates that PIF1 and HY5 reciprocally and sequentially control miR408 accumulation before and after germination (Figure 12). This circuit may form a switch for optimizing germination and the ensued post-germinative growth in light (Figure 12).

### PLC Links Copper Mobilization to Hormone Metabolism

PLC belongs to the phytocyanin family of small blue copper proteins, which are ancient copper-containing redox proteins widely distributed in microorganisms and plants (Rydén and Hunt 1993; Guss et al., 1998; De Rienzo et al. 2000; Giri et al. 2004). Different from other phytocyanins, PLCs are extremely compact with the 120-130 amino acid residues devoting to three conserved motifs. Besides the characteristic type-I copper binding motif on the C-terminus, the signal peptide and the miR408 recognition site superimpose on the N-terminus (Figure 11; Supplemental Figure 5). Our results provided two important clues to PLC function. First, through quantification of endogenous hormones, genetic analysis, gene expression profiling, and pharmacological analyses (Figure 8 to 10), we demonstrated that PLC acts as a switch downstream of PIF1 and is both necessary and sufficient to reciprocally modulate GA and ABA levels in the seed. Previously, high expression of *PLC* in the transmitting tract of the pistil was noticed (Dong et al. 2005). Over-expression of *PLC* was found to disrupt pollen tube guidance into the style and to reduce seed set (Dong et al. 2005). The latter was corroborated by observations that over-expression of miR408 resulted in larger seed size and higher grain yield (Pan et al., 2018). Thus, besides seed germination, PLC may participate in other aspects of seed biology and reproductive development. It will be interesting to test if rebalancing endogenous hormone levels is the unifying function of PLC in these processes.

Second, through RNA-sequencing, we found that *PLC* is both necessary and sufficient to regulate an overwhelming portion of the *PIF1*-dependent transcriptome underlying germination (Figure 8; Supplemental Figure 3; Supplemental Dataset 1). This finding suggest that PLC turnover is associated with major changes to cellular state. Because PLC is a vacuole located cuproprotein with a bare-bones type-I copper motif and highly expressed in mature seed (Figure 1 and 11), we contemplated that PLC is a key carrier for copper mobilization. In the seed, mineral nutrients are sequestered in the storage vacuoles (Lanquar et al., 2005; Kim et al., 2006; Roschzttardtz et al., 2009; Eroglu et al., 2017). Right after imbibition, there likely is no new assimilation of transition metals before massive translation and protein synthesis is taking place (Née et al., 2017; Paszkiewicz et al., 2017). Thus, mineral elements need to be mobilized from vacuolar stores and transported to the cytoplasm and other organelles for reconstituting biochemical activity. Disrupting these processes has been shown to lead to severe germination defects (Lanquar et al., 2005; Kim et al., 2006). Previously we have shown that miR408 promotes copper allocation to the plastid and enhances photosynthesis via elevating plastocyanin abundance (Zhang et al., 2014; Pan et al., 2018). Taken together, we speculate that the miR408-*PLC* module controls copper redistribution between the vacuole and the plastid.

### How Does PLC Regulate GA and ABA Biosynthesis?

PLC turnover as a means for copper mobilization and delivery to the plastid is consistent with previous studies on the effects of copper on plastid physiology and biochemistry. Plastid is known to be the major cellular copper sink in plants (Burkhead et al., 2009), whereby the transition metal acts as cofactor for plastocyanin in the thylakoid lumen, which is indispensable as an electron carrier in the Z-scheme of photosynthesis (Molina-Heredia et al., 2003; Weigel et al., 2003), and for the copper- and zinc-containing superoxide dismutase in the stroma, which participates in neutralizing reactive oxygen species to maintain proper redox state in the plastid (Gupta et al., 1993). Copper allocation to the plastid was shown to be critical for plastocyanin abundance and activity (Weigel et al., 2003; Zhang et al., 2014; Pan et al., 2018). Copper level was also reported to impact the number of chloroplasts per cell, thylakoid stacking, and grana size (Bernala et al., 2006). It is intriguing to note that GA biosynthesis initiates in the plastid (Sun and Kamiya 1997; Yamaguchi, 2008). Thus, regulated PLC degradation may promote copper translocation or allocation to the plastid. Copper-propelled plastid development may in turn provide the structural and biochemical niche for initiating GA biosynthesis in the seed.

Alternatively, the effect of PLC turnover on hormone rebalancing may be explained by a direct impact on ABA synthesis taking place in the cytosol. In *Arabidopsis*, AAO3 encodes an aldehyde oxidase that catalyzes the last step of ABA biosynthesis, the conversion of abscisic aldehyde to ABA (Seo et al., 2000). AAO3 is a cytosolic molybdoenzyme that requires the molybdenum cofactor for catalytic activity (Seo et al., 2000). Structural and biochemical analyses have shown that the final step of molybdenum cofactor biosynthesis is dependent on a copper-dithiolate complex, which protects the reactive dithiolate before molybdenum insertion (Kuper et al., 2004). It could be speculated that PLC is one of the copper donors, through unidentified cytoplasmic chaperones, passes on copper to the dithiolate group for synthesizing the molybdenum cofactor (Peñarrubia et al., 2015). Induction of *PLC* expression during late seed development (Figure 1) is consistent with this scenario whereby elevated PLC helps to maintain AAO3 activity and hence ABA accumulation during seed maturation. Upon light irradiation, rapid PLC turnover would deplete copper supply for AAO3 and impede ABA synthesis, which is consistent with the fourfold decline of ABA content in the *plc* seed over the wild type (Figure 10C). The finding in rice that exogenous copper increases ABA accumulation and inhibits germination (Ye et al., 2014) provides another line of evidence for this model. Further studying PLC-related copper homeostasis could shed more light on hormone synthesis and balancing during seed development and germination.

## METHODS

### Plant Materials and Growth Conditions

The wild type *Arabidopsis thaliana* used in this study was Col-0. The *pif1, PIF1-OX, pMIR408:GUS, miR408-OX*, and *amiR408* plants were as previously described (Oh et al., 2004; Zhang and Li, 2013; Zhang et al., 2014). To delete *PLC*, a CRISPR/Cas9 system employing the modified pCAMBIA1300 vector was used (Mao et al., 2013) in which the *35S* and the *AtU6-26* promoter respectively drive *Cas9* and a pair of sgRNAs that were designed to target both ends of the *PLC* coding region. The resulting construct was used to transform the wild type and T_1_ plants were individually genotyped by PCR and sequencing to identify deletion events. Approximately 100 individual T_2_ plants were genotyped to identify *Cas9*-free homozygous *plc-2* lines. The *iPLC-OX* transgenic plants were obtained by cloning the *PLC* coding sequence into the pER8 vector (Zuo et al., 2000) and transforming wild type plants. Homozygotes were selected for Hygromycin resistance in the T_2_ population. The *pPLC:PLC-GFP* transgenic plants were obtained by cloning the *PLC* coding sequence into the modified pJim19-GFP vector and substituting the *PLC* promoter for the *35S* promoter. Following transformation of the wild type plants, homozygotes were selected by Kanamycin resistance in the T_2_ population. Sequences of the relevant primers are listed in Supplemental Table 1. The *pMIR408:GUS/pif1, PIF1-OX miR408-OX, pif1 amiR408, PIF1-OX plc, lac12 lac13*, and *plc lac12 lac13* lines were generated by crossing and selection for homozygotes at the F_2_ generation.

Adult *Arabidopsis* plants were grown in soil at 22°C, ∼60% relative humidity, and under long day (16 h light/8 h dark) condition in a growth chamber. For each experiment, the seeds were harvested at approximately the same time. After harvesting, the seeds were dried at room temperature for six to eight weeks prior to germination and other experiments.

### Germination Assays

The far-red light induced germination assay was performed as described with minor modifications (Oh et al., 2004). Briefly, a triplicate set of 50-75 seeds for each sample was surface sterilized with liquid bleach and plated on half-strength MS aqueous agar medium (0.6% agar, 1% sucrose, pH 5.7). One hour after the start of sterilization, the plated seeds were irradiated with 3.2 μM m^-2^ s^-1^ far-red light for 5 min and then incubated in the dark for 48 h. For phyA_OFF_, the imbibed seeds were continuously placed in darkness for up to 120 h. For phyA_ON_, the seeds were treated with a second far-red irradiation for 4 h and then in darkness for up to 120 h. For pharmacological analysis, 100 μM paclobutrazol, 10 μM GA_3_, or 5 μM ABA (Sigma-Aldrich) was supplemented to the medium. Germination was determined by examining radicle formation at the indicated time points.

### Analysis of Seedling Greening

The seedlings were grown in dark for four days and then transferred to continuous white light (100 μM m^-2^ s^-1^) for 24 h. The greening rate was determined and calculated as the ratio of green seedlings over the total germinated seedlings as previously described (Zhong et al., 2009). Pigments were extracted from etiolated seedlings in the dark at room temperature using 90% acetone containing 0.1% NH_3_ as previously described (Zhong et al., 2014). Supernatants containing the pigments were subject to fluorescence spectral analysis using an Infinite M200 microplate reader (Tecan). The excitation wavelength was 443 nm and the emission spectra were recorded from 610 to 740 nm with 1 nm bandwidth. All measurements were performed on at least three independent biological samples and one representative set of results was shown.

### Transcript Quantification

Total RNA from imbibed seeds was isolated using the Quick RNA Isolation Kit (Huayueyang). For each experiment, mRNA and miRNA from three independent biological samples were reverse transcribed into cDNA using the PrimeScript II 1^st^ Strand cDNA Synthesis Kit (TaKaRa) and the miRcute Plus miRNA First-Stand cDNA Synthesis Kit (Tiangen), respectively. qPCR was performed using SYBR Green Mix (TaKaRa) on the 7500 Fast Real-Time PCR System (Applied Biosystems). *Actin7* and 5S RNA were used for mRNA and miRNA normalization, respectively. Sequences of the primers are listed in Supplemental Table 1.

### RNA-sequencing Analysis

Seeds of the wild type, *miR408-OX* and *pif1* were grown on half-strength MS medium. The *iPLC-OX* seed grown on medium supplemented with or without 5 μM β-estradiol (Sigma-Aldrich). All seeds were treated with the phyA_OFF_ condition for 12 h before sample collection. Total RNA was isolated using the Quick RNA Isolation Kit (Huayueyang). Library preparation and RNA-sequencing were performed on the Illumina HiSeq 2000 platform. For each genotype, three paired-end libraries from independent biological samples were prepared. At least 16 million raw paired-end reads were generated from each library. Quality control was conducted using fastQC. Cutadapt and a custom Perl script were used to trim adaptors with the parameter Q30 and the first nine bases following the adaptors with low fastQC score. After trimming, only reads longer than 100 bases were retained and the R1 and R2 files were paired simultaneously. The clean reads were mapped to the TAIR10 *Arabidopsis* genome build using STAR with an average mapping rate of ∼90% and unique mapping rate above 80%. Transcript quantification results generated by Stringtie were processed by Cuffdiff to identify differentially expressed genes. Clustering and correlation analyses were performed and visualized using R scripts. GO analysis was carried out using AgriGO (http://bioinfo.cau.edu.cn/agriGO/).

### ChIP-qPCR

ChIP was carried out on four-day-old dark-grown *PIF1-OX* and wild type seedlings on MS medium using an anti-MYC polyclonal antibody (Sigma-Aldrich) as described (Pfeiffer et al., 2014). After ChIP, equal amount of input DNA was subjected to qPCR analysis of the target DNA fragment. Fold of enrichment was calculated between *PIF1-OX* and the wild type input.

### Immunoblotting

The *pPLC:PLC-GFP* line was used for protein analysis. Dry seed was incubated in water at room temperature for 0.5 h or at 4°C for 24 h before sample collection. Total protein was isolated with an extraction buffer containing 50 mM Tris·HCl, pH 7.5, 6 mM NaCl, 1 mM MgCl_2_, 1 mM PMSF, and 1× protease inhibitor mixture (Roche). Immunoblotting was performed with an anti-GFP antibody (Abcam). An anti-RPT5 antibody (Abcam) was used as loading control.

### Histochemical Staining for GUS Activity

After stripping away the seed coat in green safe light, *pMIR408:GUS* and *pMIR408:GUS/pif1* seeds grown for 12 h in phyA_OFF_ and phyA_ON_ were incubated in a standard GUS staining solution for 3 h at 37°C. Following removal of the staining solution, seeds were washed with several changes of 75% ethanol until pigments were no longer visible. Images of the GUS staining pattern were taken with a digital camera.

### Live-cell Imaging

The *pPLC:PLC-GFP* seed was treated with phyA_OFF_ and phyA_ON_ for the indicated duration. Cotyledons were dissected away from the testa and endosperm by the application of gentle pressure to seed held between a microscope slide and a cover slip. Fluorescence images were obtained using a Nikon A1R si+ laser scanning confocal microscope equipped with an APO 40×1.25 NA water immersion objective. Excitation and emission wavelengths were 488/500 to 550 nm for GFP and 405/425 to 475 nm for vacuole autofluorescence. Autofluorescence spectra were obtained using a 32-PMT spectral detector. Spectral unmixing and image analysis were performed using the NIS Elements AR software (Nikon Instruments).

Transient expression in onion epidermal cells was performed as previously described (Wang and Frame, 2009). The gold particles were coated with plasmid DNA containing the expression cassette for PLC-GFP or COPT5-mCherry. The Biolistic PDS-1000/He Particle Delivery System (Bio-Rad) was used for bombarded with the following settings: 1,100 psi rupture disc, 25-26-inch Hg vacuum, and target distance of 10 cm. After bombardment, the explants were kept in dark at 25°C for 16-18 h and observed with the Nikon A1R si+ microscope. Excitation and emission wavelengths were 488/500 to 550 nm for GFP and 561/570 to 620 nm for mCherry.

### Quantification of Endogenous GA and ABA

For each genotype, 500 mg of seed grown in phyA_OFF_ for 24 h was collected and ground into fine powder in liquid nitrogen. Endogenous ABA was purified and measured as previously described (Fu et al. 2012) with minor modifications to the detection procedure. Briefly, UPLC-MS/MS analysis was performed on a UPLC system (Waters) coupled to the 5500 Qtrap system (AB SCIEX). Chromatography separation was achieved with a BEH C18 column (Waters) with mobile phase 0.05% HAc (A) and 0.05% HAc in ACN (B). The gradient was set initially with 20% B and increased to 70% B within 6 min. ABA was detected in the MRM mode with transition 263/153 and the isotope dilution method was used for quantification.

Quantitative GA measurement was performed as previously described using UPLC-MS/MS (Ma et al., 2015). Chromatography separation was achieved with a BEH C18 column (Waters) with mobile phase 0.05% HAc (A) and 0.05% HAc in ACN (B). Gradient was set as the following: 0-17 min, 3% B to 65% B; 17-18.5 min, 65% B to 90% B; 18.5-19.5min, 90% B; 19.5-21 min, 90% B to 3% B; and 21-22.5 min, 3% B. GA_4_ was detected in the negative MRM mode and quantified with a MRM transition. The source parameters were set as IS voltage -4500 V, TEM 600°C, GS1 45, GS2 55, and curtain gas 28.

### Transient Expression in Tobacco Protoplasts

The promoter sequence of *MIR408* was cloned from *Arabidopsis* genomic DNA, inserted into the pGreen II 0800-LUC vector (Hellens et al., 2005), and used as the reporter. The *PIF1* coding sequence was cloned from *Arabidopsis* cDNA, inserted into pGreen II 62-SK (Hellens et al., 2005), and used as the effector. Tobacco protoplasts were freshly prepared as described previously (Yoo et al., 2007). The effector or the empty vector was combined with the report construct and used to transiently transform the protoplasts using the Dual-Luciferase Reporter System (Promega), following the manufacturer’s instruction. Transfected protoplasts were incubated under low light for 16 h. The chemiluminescence was determined using a LB942 Multimode Reader (Berthold Technologies).

### Domain and Phylogenetic Analyses of PLC

PLC in *Arabidopsis* was used as query to perform a BLASTP search against all proteins in 52 plant species covering all main clades of land plants. A total of 276 PLC sequences were identified based on two criteria: E-value ≤ e^-10^ and a “plantacyanin” annotation term assigned by InterProScan. N-terminal signal peptide was predicted using SignalP (Almagro Armenteros et al., 2019). Binding site for miR408 was predicted using psRNATarget (Dai et al., 2018). To show the evolutionary trajectory, PLCs from 14 representative plant species and the most homologous genes encoding small blue copper proteins from *Physcomitrella patens, Salvinia cucullate*, and *Azolla filiculoides* were selected and mapped to a species tree obtained from TimeTree (http://www.timetree.org/).

## Supplemental Data

Supplemental Figure 1. Generation and Characterization of *PLC*-related Mutants.

Supplemental Figure 2. The miR408-*PLC* Module Specifically Regulates Germination.

Supplemental Figure 3. Enriched GO Terms Associated with *PIF1* and *PLC* Coregulated Genes.

Supplemental Figure 4. Exemplar Genes Regulated by the *PIF1-PLC* Pathway.

Supplemental Figure 5. PLC Has a Compact Domain Organization.

Supplemental Table 1. Oligonucleotide Sequences of the Primers Used in This Study.

Supplemental Dataset 1. List of *PIF1* and *PLC* Coregulated Genes.

Supplemental Dataset 2. Expression Profile of 218 Germination Related Genes.

## Accession Number

Sequence data from this article can be found in the *Arabidopsis* Genome Initiative or GenBank/EMBL databases under the following accession numbers: *MIR408* (At2g47015), *PIF1* (AT2g20180), *HY5* (At5g11206), *PLC* (At2g02850), *LAC3* (At2g30210), *LAC12* (At5g05390), *LAC13* (At5g07130), *GA2ox8* (At4g21200), *GA3ox1* (At1g15550), *GA2ox2* (At1g30040), *ABA1* (At5g67030), *NCED6* (At3g24220), *NCED9* (At1g78390), *SOM* (At1g03790), *RVE2* (At5g37260), *JMJ22* (At5g06550) and *MAN7* (At5g66460). T-DNA insertion mutants used are *pif1* (SALK_072677), *plc-1* (SALK_091945), *lac3* (SALK_031901C), *lac12* (SALK_087122), and *lac13* (SALK_023935). RNA sequencing data can be found at the National Center for Biotechnology Information Sequence Read Archive under accession number PRJNA633227.

## Author Contributions

L.L. designed and supervised the research. A.J., J.P., Y.Z., D.Z., and C.H. performed the experiments. Z.G. (Guo), Z.G. (Gao), and S.Z. analyzed the data. P.X. and J.C quantified hormone levels. A.J. and L.L. wrote the paper.

## Acknowledgements

We thank Drs. Xiangdong Fu, Xing-Wang Deng, and Hongwei Guo for providing some of the plasmids and seeds used in this study. This work was supported by grants from the National Key Research and Development Program of China (2017YFA0503800) and the National Natural Science Foundation of China (31621001).

